# Spatial proteomics on repurposed sequencers reveals early astrocytic remodeling in an ALS mouse model

**DOI:** 10.64898/2026.11.08.765207

**Authors:** Kunal Pandit, Craig Fouts, Sarah Rodwin, Silas Maniatis, Karan Dhingra, Selma Marrakchi, Daniel Domovic, Jagjit Singh, Bianca Dumitrascu, Hemali Phatnani, Maroš Pleška, Sanja Vickovic

## Abstract

Spatial proteomics can reveal how disease reshapes cells and their local environments, but extending these measurements across replicated tissue cohorts requires validated antibodies, consistent sample processing and integrated analysis. These requirements are particularly pressing in studies of amyotrophic lateral sclerosis (ALS), where pathology evolves unevenly across spinal-cord regions and disease stages. Here, we established an end-to-end spatial proteomics workflow linking a validated 28-antibody panel to automated fluorescence decoding and imaging on repurposed high-throughput sequencers, followed by multicycle image processing, cellular profiling and probabilistic spatial-neighborhood analysis. We applied it to 70 lumbar spinal-cord sections from nine SOD1-G93A and nine wild-type mice at presymptomatic, disease-onset and symptomatic stages. Spatial neighborhood analysis revealed that the environments surrounding motor neurons were already remodeled at disease onset. GFAP-high neighborhoods expanded in the ventral gray matter, with 43% of motor neurons embedded in these environments in SOD1-G93A mice, compared with 15% in wild-type mice. This early expansion preceded the later decline of non-reactive gray matter and detectable motor-neuron loss. The regional organization of the protein-defined changes shared features with neuronal and astrocytic patterns mapped by spatial transcriptomics in mouse and human ALS. This work enables spatial proteomics across replicated tissue cohorts and identifies localized remodeling of the environments around surviving motor neurons as an early feature of disease in the SOD1-G93A model.

## INTRODUCTION

Tissue function emerges from spatially organized communities of cells whose identities, molecular states and interactions define local microenvironments^1,2^. Disease progression can alter both individual cells and the organization of their surrounding tissue, producing spatial patterns that evolve across disease stages^3–5^. Such spatial organization is particularly relevant in amyotrophic lateral sclerosis (ALS), where interactions with surrounding astrocytes, microglia, oligodendrocytes and immune cells shape the regional and temporal progression of motor-neuron degeneration^5–9^. Understanding ALS progression therefore requires following both motor neurons and the local environments they occupy.

Spatially resolved studies in experimental models and human post-mortem tissue have begun to map ALS pathology within its native tissue architecture. In SOD1-G93A mice, a widely used model of ALS, spatial transcriptomics identified region- and stage-specific gene-expression programs, including early astrocyte- and microglia-associated changes^5^. In human ALS spinal cord, spatial profiling has revealed immune and myeloid programs associated with established pathology^10^ and loss of motor-neuron and synaptic programs in the ventral horn^11^. These studies establish that ALS pathology is spatially structured, but how the protein-defined environments surrounding motor neurons change across disease stages remains less clear. A central question is whether these environments are already remodeled while motor neurons are still present. Addressing this question requires cellular-scale spatial proteomics across biological replicates and disease stages, linking changes in the surrounding tissue to the molecular state and abundance of motor neurons.

High-resolution imaging-based spatial proteomics can measure protein-defined states and neighborhoods while preserving cellular morphology and tissue architecture^3,4,12^. Applying it across replicated tissue cohorts, however, requires coordination of molecular detection, instrumentation and analysis. Cyclic immunofluorescence^13^ and DNA-barcoded antibody methods^14,15^ overcome the multiplexing limits of conventional fluorescence imaging through successive rounds of molecular readout. Among these methods, ImmunoSABER uses DNA concatemers to amplify antibody-associated signals without repeated antibody staining. These workflows require repeated rounds of reagent exchange and imaging, making them laborious to perform manually. Dedicated commercial systems reduce this burden by automating both steps and thus support the consistent execution of established workflows^16^. Developing or adapting novel assays, however, requires a degree of flexibility in reagent exchange, cycle timing, and image acquisition that commercial platforms may not offer^17^. This limitation highlights the need for programmable platforms that support both assay development and consistent execution across tissue cohorts.

To make automated imaging more flexible and accessible, we previously developed PySeq2500, an open-source toolkit that repurposes decommissioned Illumina HiSeq 2500 sequencers for programmable fluorescence imaging^18^. Here, we combine this platform with a validated 28-antibody ImmunoSABER panel and two new analysis tools to establish an integrated spatial proteomics workflow. First, Hudson converts multicycle images into spatially resolved cellular-segment profiles. Second, sceLDA identifies recurring tissue neighborhoods from their molecular features and spatial context. We apply this newly developed end-to-end workflow to 70 lumbar spinal-cord sections from nine SOD1-G93A and nine wild-type mice at presymptomatic, disease-onset and symptomatic stages. At disease onset, motor neurons show altered protein signals, and GFAP-high neighborhoods expand around them before a reduction in motor-neuron counts is detected at the symptomatic stage. These findings reveal how the local environments occupied by surviving motor neurons change as disease progresses.

## RESULTS

### Repurposed sequencers enable spatial proteomics at scale

To profile the cellular organization and molecular features of ALS pathology, we assembled a candidate panel of 30 antibodies targeting the major cellular populations and tissue compartments of mouse spinal cord. The candidate panel included markers of neurons, astrocytes, myeloid cells, oligodendrocytes, blood vessels and T cells, together with SOD1 and proteins associated with immune signaling, proliferation, cell adhesion, mitochondrial function, endolysosomal activity and lipid handling (**Fig. 1a; Supplementary Table 1**). Two non-antibody stains, DRAQ5 for nuclei and Cy3-labeled poly(dT) probes targeting cytoplasmic mRNA, were included to aid cellular segmentation (**Fig. 1a, right**). The panel was designed for ImmunoSABER, which enables multiplexed fluorescence detection using DNA-barcoded antibodies^15^ (**Fig. 1b**). Each DNA-conjugated antibody was first screened by conventional immunofluorescence using fluorescently labeled secondary antibodies. Candidates were assessed for specific signal above background and agreement with the expected anatomical distribution. The conjugated ChAT and SMI-32 candidates did not yield sufficiently specific staining under these conditions and were excluded (**Fig. 1a**). The remaining 28 antibodies resolved complementary features of lumbar spinal cord architecture across tissue and cellular scales and were retained (**Supplementary Fig. 1a,b**).

**Figure 1:**
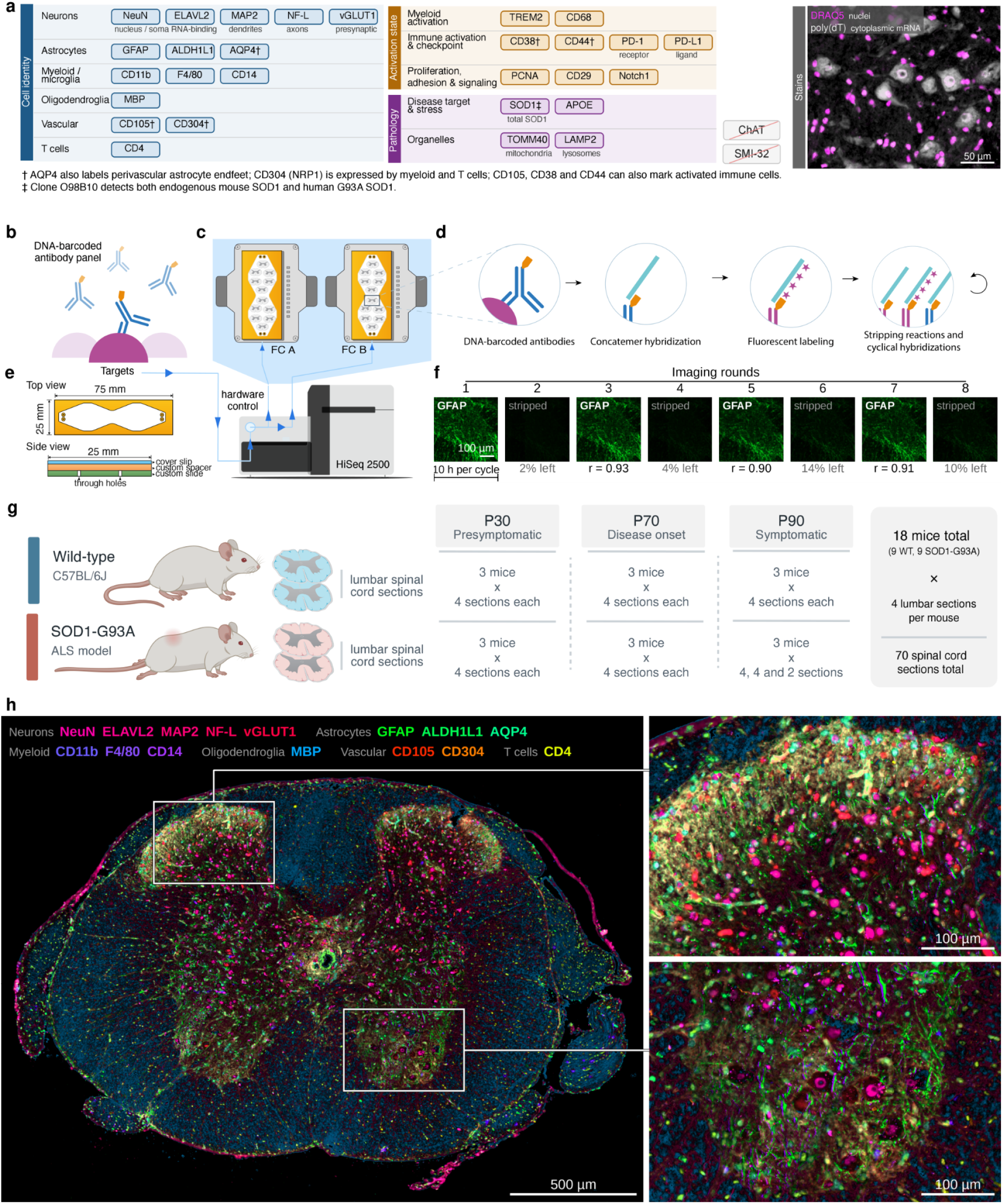
Antibody panel, automated imaging workflow and multiplexed spinal-cord images. **(a) Antibody panel and stains:** The 28 antibodies grouped by primary biological application. Two additional candidates, ChAT and SMI-32 (struck through), were excluded following screening. Right, the two stains, DRAQ5 (nuclei, magenta) and poly(dT) (cytoplasmic mRNA, gray), in the ventral horn of the section shown in h; motor neurons are distinguished by their large, mRNA-rich cytoplasm. Scale bar, 50 µm. **(b) DNA-barcoded antibody panel:** Schematic of antibodies conjugated to DNA oligonucleotides for multiplexed protein detection. **(c) Automated imaging platform:** Schematic of the Illumina HiSeq 2500 repurposed for fluorescence imaging. The instrument provides programmable control of the fluidics, imaging hardware and two flow-cell positions (FC A and FC B). **(d) Automated ImmunoSABER workflow:** Following antibody staining, DNA concatemers are hybridized to antibody-associated oligonucleotides. Successive imaging cycles of fluorescent imager hybridization, imaging and stripping enable sequential detection of multiple protein targets. **(e) Custom flow-cell design:** Top and side views of the flow cell, consisting of a custom glass slide, spacer and coverslip, with through-holes for reagent exchange. The 7.75 cm² imaging area holds up to 12 spinal cord sections. **(f) Repeatability of cyclic fluorescence imaging:** One 450 × 450 µm field of a mouse spinal cord section imaged over eight successive rounds, with the GFAP imager hybridized in odd rounds and stripped in even rounds. All images are shown with identical display settings. Values below the images were measured across the whole section: pixel-wise Pearson’s r relative to round 1 for GFAP rounds, and mean residual signal as a percentage of the preceding GFAP round for stripped rounds. One imaging cycle of a full flow cell takes about 10 h. Scale bar, 100 µm. **(g) Experimental cohort:** Study design comprising nine SOD1-G93A and nine wild-type mice at three ages (P30, P70 and P90), with three mice per genotype and age group. Four lumbar spinal cord sections were analyzed per mouse (two for one P90 SOD1-G93A mouse), 70 sections in total. **(h) Multiplexed spatial proteomic imaging:** Composite image of 15 cell-identity markers in a lumbar spinal cord section from a P90 SOD1-G93A mouse, with the dorsal horn (top) and ventral horn (bottom) regions in the white boxes enlarged on the right. Marker colors are indicated at the top of the panel; the remaining 13 markers are shown in **Supplementary Fig. 1f,g.** Scale bars, 500 µm (whole section) and 100 µm (enlarged regions).

The fluorescent readout of the 28-antibody panel was automated on repurposed HiSeq 2500 instruments, which integrate fluidics, thermal control and fluorescence imaging across two flow cells in parallel (**Fig. 1c**). Building on our previously developed PySeq2500 control software^18^, we programmed the instrument for concatemer hybridization followed by successive imaging cycles to detect individual protein targets (**Fig. 1d**). A custom-designed flow cell with 7.75 cm² of usable imaging area accommodated up to 12 mouse spinal-cord sections, allowing us to image 24 sections in a single instrument run under the same automated readout protocol (**Fig. 1e**). Across successive imaging cycles, repeated fluorescence readouts showed high repeatability (pixel-wise Pearson’s r = 0.90–0.98), while imager removal reduced fluorescence to background levels, indicating limited carryover between imaging cycles (**Fig. 1f**). Repeated detection of GFAP, CD68, MBP, TOMM40, CD29 and NeuN further demonstrated that distinct antibody targets could be reliably recovered across successive imaging cycles (**Supplementary Fig. 1c,d**). Compared with manual processing, automated imaging showed greater fluorescence-intensity repeatability across successive imaging cycles and anatomical regions (**Supplementary Fig. 1e**).

The combination of automated cycling and large-area imaging allowed us to apply the 28-antibody workflow to a cohort of 18 mice comprising nine SOD1-G93A animals and nine age-matched wild-type C57BL/6J controls. Three animals per genotype were examined at P30, P70 and P90, corresponding to presymptomatic, disease-onset and symptomatic stages, respectively. Four lumbar spinal cord sections were imaged per animal (two for one P90 SOD1-G93A mouse), yielding 70 sections in total (**Fig. 1g; Supplementary Table 3**). Composite images of 15 cell-identity markers acquired over successive imaging cycles preserved the anatomical organization of gray and white matter while revealing heterogeneous protein distributions within individual spinal cord sections (**Fig. 1h; Supplementary Fig. 1f,g**). At cellular resolution, the dorsal horn contained densely packed small neurons, whereas in the ventral horn, large motor-neuron somata lay among astrocytic processes (**Fig. 1h**). Together, automated cycling and large-area imaging yielded matched 28-protein maps from 18 mice at three disease stages, with which we next examined motor neurons and their neighborhoods.

### Motor-neuron protein signals and surrounding GFAP are altered at disease onset

To extract quantitative information from the multiplexed fluorescence images, we developed Hudson, an integrated image-analysis pipeline that performs background correction, channel registration, spectral unmixing, cellular segmentation and feature extraction (**Supplementary Fig. 2a**, **Methods**). A cellular segmentation model refined through human-in-the-loop training improved the delineation of cellular structures compared with the original model (**Supplementary Fig. 2b,c**). Across 70 spinal cord sections, Hudson generated 805,334 spatially resolved segments containing protein intensities, morphological measurements and vision transformer (ViT) image embeddings (**Supplementary Fig. 2d**). The segmented objects did not always correspond to individual cells: many encompassed cell bodies, whereas others represented disconnected cellular processes or structures overlapping neighboring cells (**Supplementary Fig. 2e**). We therefore refer to these analytical units as cellular segments rather than assuming a one-to-one correspondence with individual cells.

We first asked whether the workflow captured expected motor-neuron pathology and detected changes before neuronal loss. To this end, we examined motor neurons, whose progressive degeneration is a hallmark of the SOD1-G93A model^19^ (**Fig. 2a**). Motor neurons were annotated manually by their ventral-horn location, large soma morphology and neuronal-marker staining (**Fig. 2b**). Their abundance was comparable between SOD1-G93A and wild-type mice at P30 and P70, but was reduced by about one third in SOD1-G93A mice at P90 (0.65-fold; 95% CI 0.51–0.83; **Fig. 2c**). This reduction was directly observable as depletion of large neuronal somata from the ventral horn in representative images (**Fig. 2b**). To examine whether molecular changes preceded neuronal loss, we next compared marker signals in motor-neuron segments between genotypes. Each marker signal was expressed relative to all cell-sized, non-motor-neuron segments inside the cord in the same section, which removes section-wide differences in staining and imaging (**Fig. 2d**). No marker differed at P30. At P70, when motor-neuron numbers were still unchanged, nominal decreases (p < 0.05) were observed in several motor-neuron signals, including the neuronal marker NeuN, the adhesion receptor integrin β1 (CD29), and SOD1, the antioxidant enzyme whose mutant human form drives disease in this model. Motor-neuron segments also showed a higher relative signal of the astrocytic marker GFAP (**Fig. 2d**).

**Figure 2:**
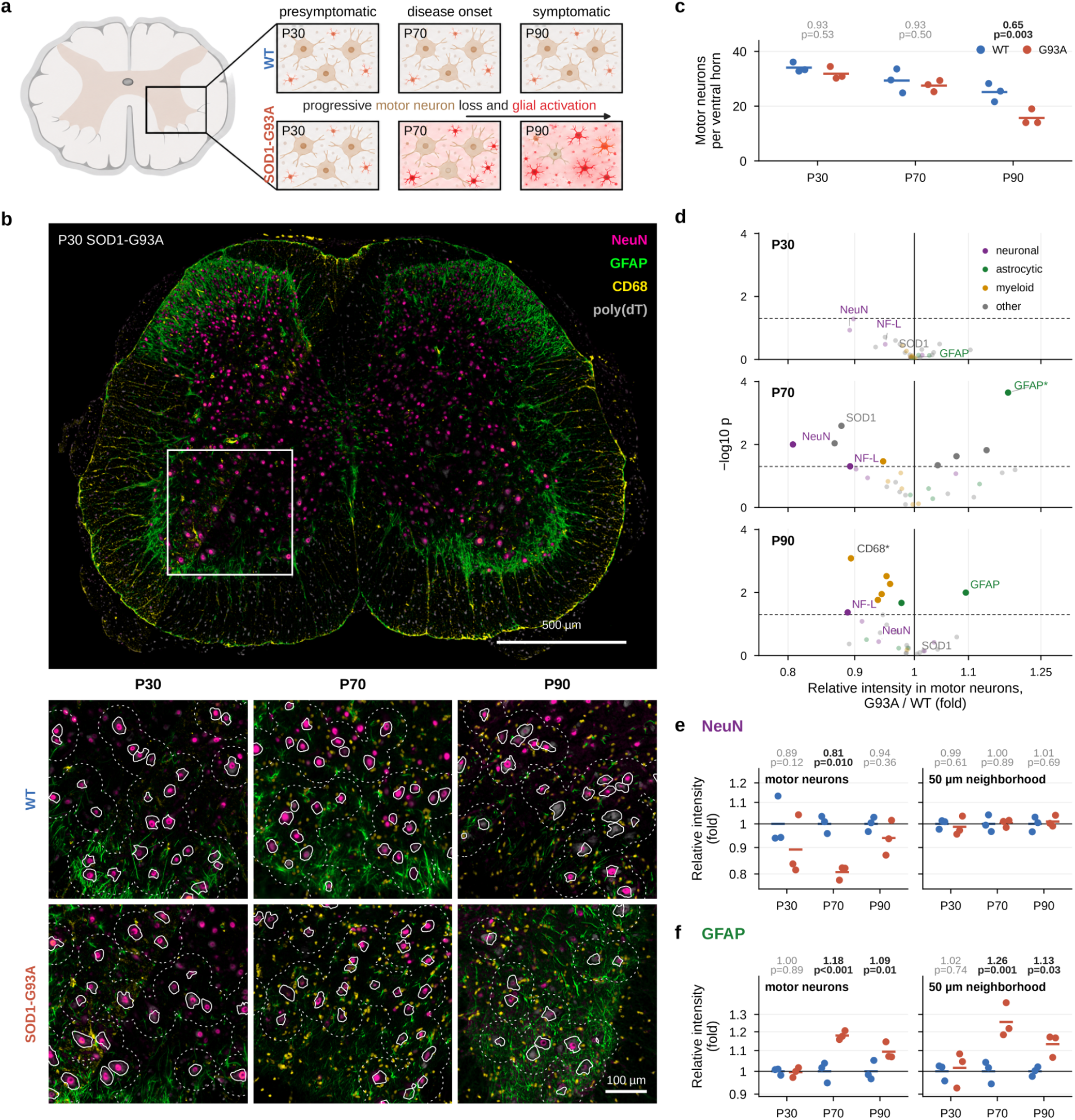
Motor-neuron counts and protein signals across disease stages. **(a) Study overview:** Schematic of the lumbar spinal cord and ventral-horn changes in wild-type and SOD1-G93A mice at P30 (presymptomatic), P70 (disease onset) and P90 (symptomatic). **(b) Motor neurons in the ventral horn:** Composite image of NeuN (magenta), GFAP (green), CD68 (yellow) and poly(dT) (gray) in a lumbar spinal cord section from a P30 SOD1-G93A mouse (top), with one ventral horn per genotype and age enlarged below. White outlines mark the segmented motor neurons, and dotted lines mark 50 µm from motor-neuron centers (the neighborhoods analyzed in e and f). Wild-type and SOD1-G93A images of each age are displayed with identical settings. Scale bars, 500 µm (top) and 100 µm (enlarged regions). **(c) Motor-neuron counts:** Motor neurons per ventral horn. Dots, mice; lines, group means. **(d) Protein changes in motor neurons:** Volcano plots of all 30 imaged channels (28 proteins, the DRAQ5 nuclear stain and poly(dT)) in motor-neuron segments, SOD1-G93A vs wild type at each age. Relative intensities in d–f were normalized to the median signal of all cell-sized, non-motor-neuron segments inside the same spinal cord section. Colors indicate the cell types that predominantly express each marker; faint points, p ≥ 0.05; dashed line, p = 0.05. Asterisks indicate Benjamini–Hochberg q < 0.05 across 90 tests (30 channels × 3 ages). Signals from astrocytic and myeloid markers in motor-neuron segments can include contributions from neighboring cellular processes. **(e) NeuN intensity:** Relative NeuN intensity in motor-neuron segments and in their 50-µm neighborhoods (cell-sized, non-motor-neuron segments within 50 µm of a motor-neuron center), shown as fold changes relative to age-matched wild-type mice. Dots, mice; lines, group means. **(f) GFAP intensity:** As in e, for GFAP. In c, e and f, numbers above the plots give the SOD1-G93A/wild-type ratio and two-sided p value. Motor-neuron counts comprised 3,514 annotated motor-neuron profiles across 127 ventral horns in 69 sections from 18 mice. Of these profiles, 3,451 were matched to cellular segments for the protein-signal analyses in d–f. Dots represent mice; measurements from the available sections were combined within each mouse. n = 3 mice per genotype and age, with mice as the unit of analysis (Methods). G93A, SOD1-G93A.

Because motor-neuron segments can include signals from neighboring cellular processes, we further compared NeuN and GFAP in motor-neuron segments and in surrounding segments within 50 µm of their centers (**Fig. 2b, dotted outlines**). Relative NeuN signal was lower in SOD1-G93A motor-neuron segments at P70 (0.81-fold) but did not differ detectably in their 50-µm neighborhoods (**Fig. 2e**). In contrast, relative GFAP signal was higher in SOD1-G93A mice in both motor-neuron segments and surrounding non-motor-neuron segments at both P70 (1.18- and 1.26-fold) and P90 (1.09- and 1.13-fold; **Fig. 2f**). Because motor neurons do not express GFAP, the higher signal within motor-neuron segments likely reflects an increased presence of astrocytic processes surrounding or contacting the neurons. Together, these analyses recovered the expected progression of motor-neuron pathology, with reduced relative NeuN signal at P70 preceding the loss of about one third of motor neurons by P90. The accompanying GFAP enrichment around surviving neurons prompted us to examine how these altered local environments were organized across the ventral horn and how their spatial extent changed with disease.

### Spatial topic modeling reveals reorganization of tissue neighborhoods in the symptomatic ventral horn

Characterizing tissue neighborhoods in the spinal cord required identifying recurring local mixtures of cellular phenotypes from protein intensities, morphology and image features. Importantly, clustering cellular profiles without spatial context does not directly capture which phenotypes occur together in tissue neighborhoods. We therefore developed single-cell embedded latent Dirichlet allocation (sceLDA), which combines a learned representation of these diverse features with spatial topic modeling to identify neighborhoods that can overlap and vary in composition (**Fig. 3a, Methods**). The approach draws on topic modeling, which represents documents as mixtures of recurring topics^20,21^. In sceLDA, morphological features, protein intensities and ViT image embeddings are first encoded as latent embeddings using a variational autoencoder (VAE), spatially smoothed and converted into discrete phenotypic words. Spatial latent Dirichlet allocation (SLDA) then identifies recurring combinations of these words and maps their distribution across the tissue (**Supplementary Fig. 3a**).

**Figure 3:**
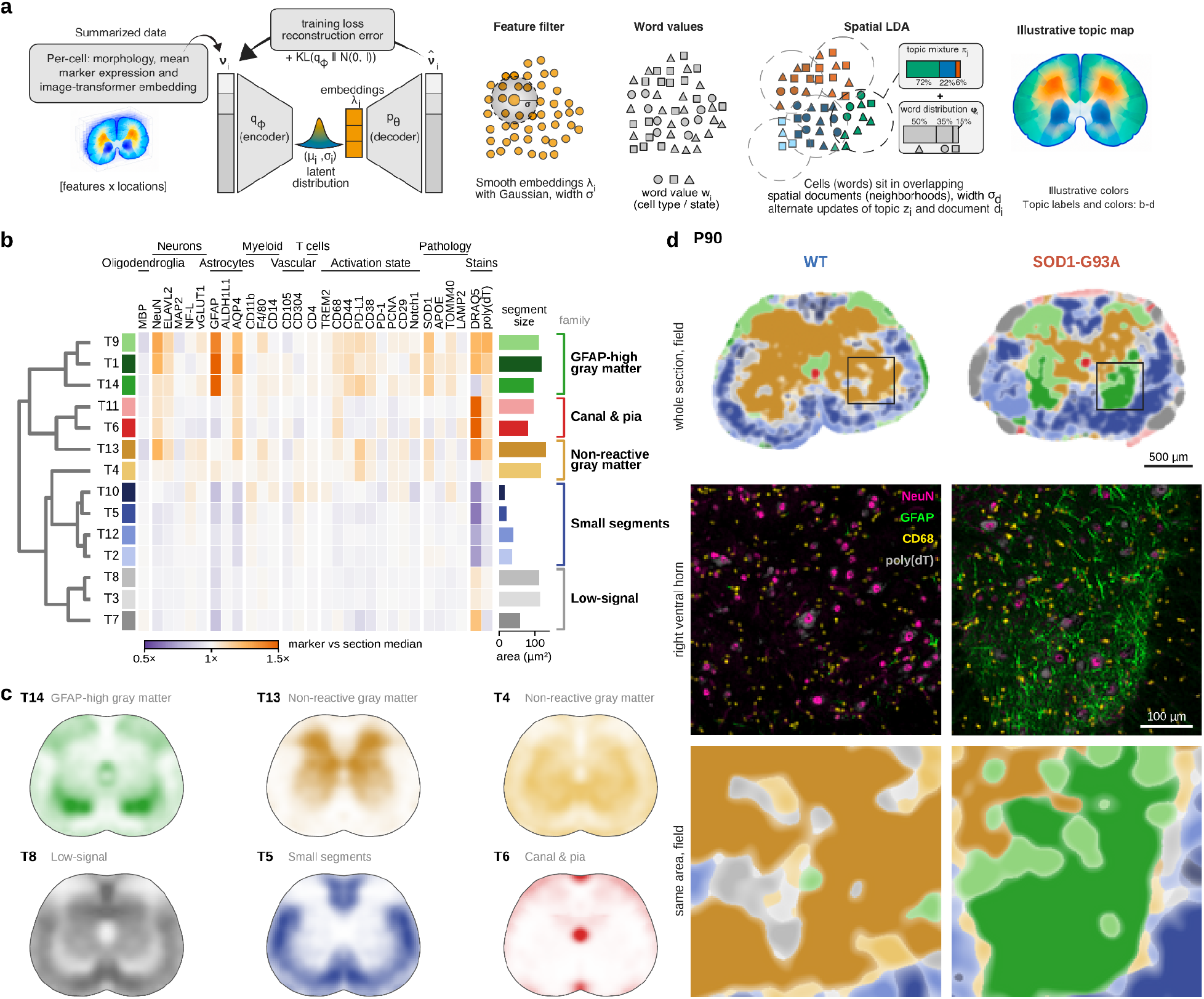
Spatial topic modeling and tissue-neighborhood maps of the mouse spinal cord. **(a) Schematic of sceLDA:** Single-cell embedded latent Dirichlet allocation (sceLDA) combines a variational autoencoder (VAE) with spatial latent Dirichlet allocation (SLDA). Morphological features, protein intensities and ViT image embeddings of each segment are encoded as VAE latent embeddings, spatially smoothed and discretized into phenotypic word values. SLDA then identifies recurring topics from the local distributions of these words, producing spatial maps of topic membership. Colors in a are illustrative; the 14 topics and their family colors are shown in b–d. **(b) Marker profiles of the topics:** Median intensity of each marker in the segments of each of the 14 topics, relative to the median of all segments in the same section, averaged over sections and then over mice (n = 18). Bars give the median segment area. The tree groups topics by how much their marker profiles differ relative to the variation between mice (complete linkage); colors mark the resulting families, with one shade per topic. **(c) Topic locations:** Share of segments assigned to one representative topic of each family at each position of the spinal cord. All 70 sections were registered to a common outline by anatomical landmarks, and left and right sides, sections and mice were averaged. Each map is scaled to its own maximum. **(d) Neighborhoods at P90:** Wild-type and SOD1-G93A sections imaged on the same slide. Top, whole sections colored by the most frequent topic among neighboring segments, paler where topics mix (colors as in b); boxes mark the enlarged regions. Middle, right ventral horn stained for NeuN (magenta), GFAP (green), CD68 (yellow) and poly(dT) (gray), displayed with identical settings. Bottom, the same regions colored as in the top row. Scale bars, 500 µm (top) and 100 µm (middle and bottom).

To test whether this approach improved neighborhood identification over existing methods, we compared sceLDA with spatially aware methods (SLDA, MEFISTO, NSF and NSFH) and non-spatial methods (LDA, NMF and k-means) on synthetic datasets with known spatial topics. Across 10 synthetic datasets, sceLDA achieved higher median pixel-wise accuracy, weighted Jaccard index and Calinski–Harabasz index than the compared methods (**Supplementary Fig. 3b–e**). Separate runtime measurements showed computation time comparable to SLDA (**Supplementary Fig. 3f**). Applied to the spatially resolved profiles generated by Hudson, sceLDA identified 14 topics with distinct marker profiles and anatomical distributions, which fell into five families (**Fig. 3b,c**): GFAP-high gray matter, including a dorsal-horn topic (T1) and a ventral-horn topic (T14); non-reactive gray matter (T4 and T13); the central canal and pia; and two families defined by small segment size or by uniformly low marker signals.

We first examined the fully developed reorganization at the symptomatic stage, then asked when GFAP-high neighborhoods began to expand. At P90, the neighborhood maps revealed a marked reorganization of the ventral horn (**Fig. 3d**). In wild-type mice, the ventral gray matter was dominated by the non-reactive gray-matter topic T13, enriched in the neuronal markers NeuN and ELAVL2 without elevated GFAP (**Fig. 3b**). In SOD1-G93A mice, this neighborhood was largely replaced by two GFAP-high topics: T14, defined by the reactive-astrocyte markers GFAP and CD44^22^, and T9, which combined neuronal and astrocytic features and was found mainly in dorsal regions in wild-type mice. The remaining motor neurons were thus embedded in astrocyte-rich environments, in agreement with the dense GFAP-positive processes visible in the corresponding images (**Fig. 3d**). This pattern was consistent with the reactive astrogliosis described in this model^23^. The expansion of T9 into the ventral horn captured neighborhoods that retained neuronal features while becoming enriched in astrocytic signals. Representing these features together allowed us to characterize changes around surviving neurons without reducing their local environments to a single cell type. Together, these results showed that sceLDA recovered anatomically interpretable tissue neighborhoods and resolved their remodeling at the symptomatic stage.

### GFAP-high neighborhoods expand around motor neurons before detectable motor-neuron loss

To determine when the ventral-horn reorganization (**Fig. 3d**) began and whether it was concentrated around motor neurons, we linked the sceLDA neighborhood maps to the annotated motor-neuron positions in images acquired on decommissioned sequencers using PySeq2500 (**Fig. 4a**). We considered a motor neuron embedded in a GFAP-high neighborhood when the two GFAP-high topics of the ventral horn, T14 and T9, predominated among the cells within a 50 µm radius. At P30, the two genotypes did not differ. By P70, when motor-neuron numbers were still unchanged but their protein profiles had already shifted (**Fig. 2c–e**), 43% of motor neurons in SOD1-G93A mice were embedded in GFAP-high topics, compared with 15% in wild-type mice. At P90, the proportions were 53% and 6% (**Fig. 4a,b**). The difference between genotypes was largest next to the motor neurons and narrowed with distance, remaining significant up to 150 µm at P70 and throughout the 200 µm examined at P90 (**Fig. 4c**). Across the ventral horn as a whole, GFAP-high neighborhoods more than doubled at P70 and remained expanded at P90, whereas non-reactive gray matter (T4 and T13) declined only at P90, by more than a third (**Fig. 4d**). These changes were largely confined to gray matter. In white matter, neither family differed between genotypes (**Fig. 4d**). The earliest detectable change was therefore an expansion of GFAP-high topics around motor neurons at P70.

**Figure 4:**
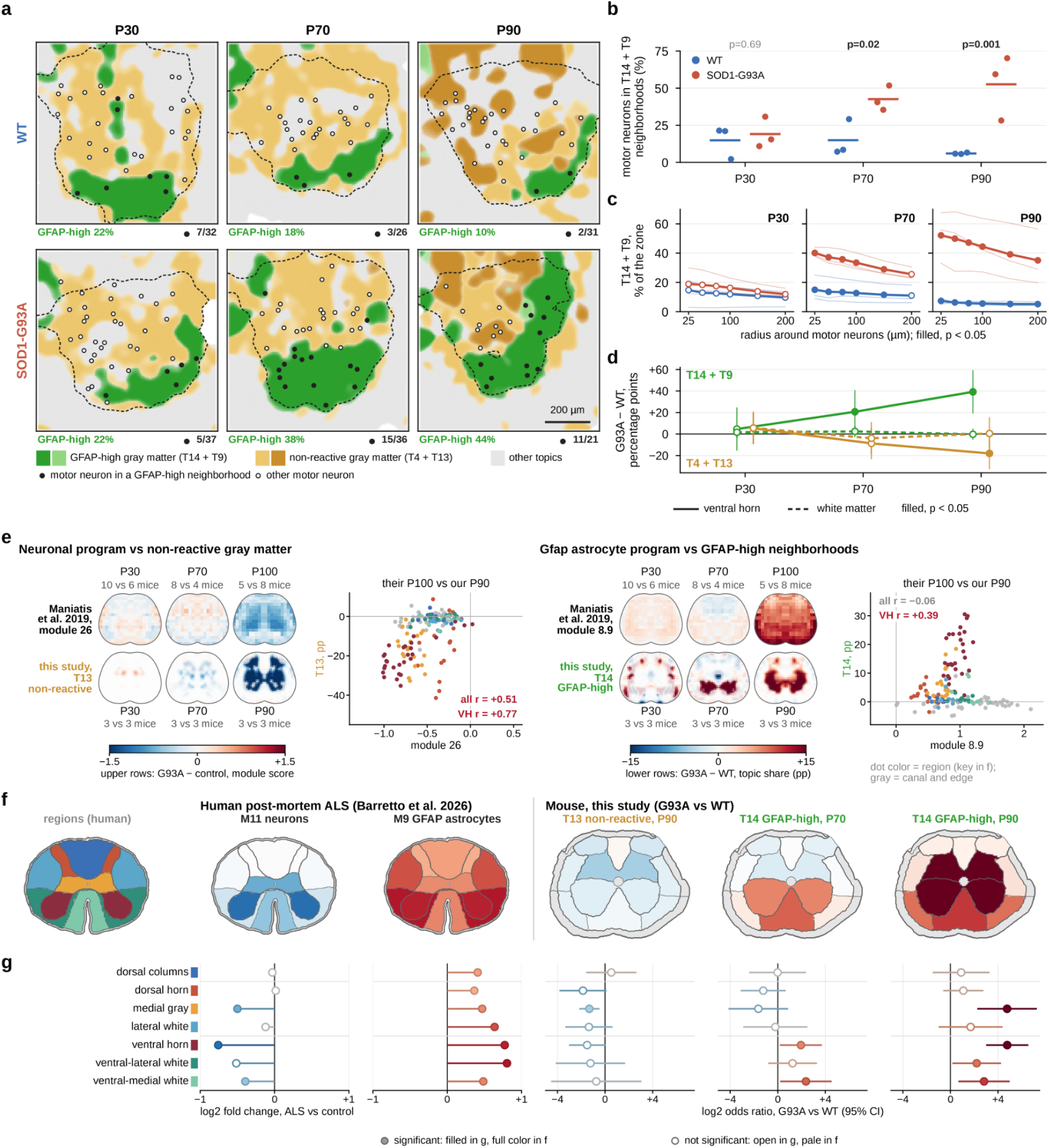
Tissue-neighborhood measurements across disease stages and comparisons with mouse and human spatial transcriptomics. **(a) Ventral-horn neighborhoods:** One ventral horn per genotype and age, colored by the most frequent topic among neighboring cell-sized segments, paler where topics mix. GFAP-high gray matter (T14 and T9, greens) and non-reactive gray matter (T13 and T4, brown and yellow) are highlighted; all other topics are gray (colors as in Fig. 3). Dashed lines mark the annotated ventral horn. Motor neurons are marked with filled circles when T14 or T9 is the most frequent topic among the cell-sized segments within 50 µm (as counted in b) and open otherwise. Percentages give the share of ventral-horn segments assigned to T14 + T9. Counts give motor neurons in GFAP-high neighborhoods divided by all annotated motor neurons. Scale bar, 200 µm. **(b) Motor neurons in GFAP-high neighborhoods:** Percentage of annotated motor neurons whose neighborhood within 50 µm has T14 or T9 as its most frequent topic. Dots, mice; lines, group means; numbers, p values for SOD1-G93A vs wild type. **(c) GFAP-high tissue around motor neurons:** T14 and T9 as a percentage of the cell-sized segments within a given distance of any motor neuron, for zones from 25 to 200 µm. Thick lines, group means (blue, wild type; red, SOD1-G93A); thin lines, mice. Filled circles, p < 0.05 for SOD1-G93A vs wild type; open circles, not significant. **(d) Ventral horn and white matter:** Difference between SOD1-G93A and wild type in the share of segments assigned to GFAP-high (T14 and T9, green) and non-reactive (T4 and T13, brown) topics, in the ventral horn (solid lines) and in white matter (dashed lines), in percentage points with 95% confidence intervals. Filled circles, p < 0.05; open circles, not significant. **(e) Comparison with mouse spatial transcriptomics:** Each block pairs a program of Maniatis et al.^5^ with the corresponding topic. Numbers above and below the maps give the mice per group, SOD1-G93A vs control. Upper rows, change in module score between SOD1-G93A and SOD1-WT mice for the neuronal gray-matter module 26 (left) and the Gfap reactive-astrocyte submodule 8.9 (right), digitized from their supplementary figures. Lower rows, change in T13 (left) and T14 (right) share (SOD1-G93A minus wild type, percentage points) on the common outline of Fig. 3c, in full color where all three SOD1-G93A mice lie above or below all three wild-type mice. Scatters, the two studies compared position by position (their P100 vs our P90), with dots colored by region as in f; r, Pearson correlation over all positions and over ventral-horn positions (VH). **(f) Comparison with human ALS spinal cord:** Left, the seven regions compared, on a schematic human lumbar spinal cord. Middle, change in module score between ALS and control donors for the neuronal module M11 and the GFAP-astrocyte module M9 in post-mortem lumbar spinal cord^11^. Right, change in T13 share at P90 and in T14 share at P70 and P90 in this study (log2 odds ratio). Red, higher and blue, lower in ALS or SOD1-G93A; pale, not significant. **(g) Regional values:** The values in f by region. Human, log2 fold change, filled where significant in Barretto et al.^11^; mouse, log2 odds ratio with 95% confidence interval, filled where p < 0.05. The motor-neuron-neighborhood analyses included 3,451 matched motor-neuron profiles across 69 sections from 18 mice. Measurements were summarized per mouse, with n = 3 mice per genotype and age.

The neighborhood changes we observed largely agreed in location with those detected by lower-resolution spatial transcriptomics in the same model (**Fig. 4e**)^5^. Briefly, the loss of non-reactive gray matter at P90 spatially coincided with the loss of a neuronal gray-matter program at P100, the symptomatic stage examined by Maniatis et al.^5^. The reactive-astrocyte program identified in the same study also increased in the ventral horn, but it rose throughout the spinal cord and only at P100. By comparison, the GFAP-high neighborhood studied here was confined to the ventral gray matter and had already expanded at P70. The P70 increase in ventral-horn GFAP remained detectable after pooling segment-level measurements into 55- and 100-µm spots (**Supplementary Fig. 4**), indicating that this regional protein signal was preserved after spatial pooling.

To assess whether the remodeling observed in SOD1-G93A mice had a regional counterpart in human ALS, we compared our neighborhood maps with spatial transcriptomic patterns in post-mortem human lumbar spinal cord^11^ (**Fig. 4f,g**). Human ALS tissue showed reduced neuronal-program scores and increased GFAP-astrocyte-program scores, most prominently in and around the ventral horn. This pattern resembled the reduction in non-reactive gray-matter topics and expansion of GFAP-high neighborhoods in symptomatic mice. The correspondence was more limited in white matter: the human astrocyte program increased throughout, whereas the mouse GFAP-high neighborhoods extended only into ventral white matter adjacent to the ventral horn. Together, these comparisons connect the regional disease programs identified by spatial transcriptomics with the protein-defined environments occupied by individual motor neurons. Spatial proteomics thus adds a local view of ALS pathology across disease stages, revealing that astrocytic remodeling is already established around surviving motor neurons at disease onset, before detectable neuronal loss.

## DISCUSSION

Spatial proteomics can connect molecular changes with the cellular organization in which disease develops, but its broader application depends on access to automated imaging systems that can be adapted to different experimental needs. We sought to make these measurements more affordable and customizable by building on PySeq2500 to repurpose decommissioned sequencing instruments for programmable protein imaging^18^. Here, we combined automated ImmunoSABER readout with Hudson for cellular-segment profiling and sceLDA for spatial-neighborhood analysis^15^. This integration allowed us to examine motor-neuron protein signals, abundance and surrounding tissue composition within the same tissue series. Applied across disease stages in SOD1-G93A mice, it revealed that localized astrocytic remodeling was already established around surviving motor neurons before detectable neuronal loss.

The workflow recapitulated established ALS pathology and identified additional changes in tissue organization. The principal biological finding was localized astrocytic remodeling at disease onset, before a detectable reduction in motor-neuron counts. At P70, GFAP-high neighborhoods expanded in the ventral horn of SOD1-G93A mice and contained about half of the annotated motor neurons, compared with one in seven in controls. In contrast, reduced motor-neuron counts were detected only at P90. Neighborhood analysis therefore captured a tissue change that was not apparent from neuronal abundance alone. This timing does not imply that astrocytic remodeling precedes neuronal injury. In SOD1-G93A mice, neuromuscular denervation and axonal degeneration can occur before motor-neuron cell bodies are lost^24^. Molecular profiling has also identified a disease-associated motor-neuron state before cell death^25^. The P70 phenotype is thus consistent with remodeling around already affected neurons, but does not establish that astrocytes initiate the disease.

Nevertheless, glial remodeling could influence disease progression even if it begins as a response to neuronal injury. In conditional SOD1-G37R mice, reducing mutant SOD1 expression in astrocytes or microglia/macrophages slowed later disease progression^6,7^. In addition, astrocytes can affect motor-neuron function without causing neuronal loss. For example, loss of astrocytic Kir4.1 altered fast α-motor-neuron properties and reduced peak muscle strength without detectable motor-neuron loss^26^. These findings support the possibility that GFAP-high neighborhoods mark changes in local neuronal support, rather than simply a response to cell loss. Neighborhood expansion could reflect altered protein expression, changes in astrocytic processes or shifts in cell abundance. Functional measurements are therefore needed to determine whether these environments preserve neuronal function, contribute to injury or combine both effects.

The regional organization of these changes complements previous spatial descriptions of ALS pathology. Spatial transcriptomics identified region- and stage-dependent neuronal and glial programs in SOD1-G93A mice^5^. More recent spatial profiling localized reactive microglia/macrophages near motor neurons early in disease^25^. Our analysis adds a protein-level view of these local environments. Human spatial transcriptomics likewise identified reduced neuronal and synaptic programs in the ventral horn, together with regional changes in astrocyte-associated programs^11^. These findings place the neighborhoods described here within a broader pattern of spatially organized neuronal and glial pathology. However, the studies differ in molecular readouts, sampling and control groups, and the human data represent end-stage disease. Regional agreement therefore supports the relevance of ventral-horn remodeling beyond this model, but does not establish an equivalent temporal sequence or mechanism across species.

Together, these findings extend the description of early ALS pathology from motor-neuron protein changes to spatially organized remodeling of the surrounding tissue. Linking these neighborhood maps to neuronal subtype, innervation and function could help determine how local environments relate to vulnerability or compensation. By combining programmable imaging on repurposed sequencers with cellular profiling and neighborhood analysis, the workflow provides a customizable way to compare these environments across disease models and measure how they respond to experimental perturbations.

## MATERIALS AND METHODS

### Animals and tissue collection

Wild-type C57BL/6J mice and hSOD1-G93A transgenic mice were obtained from The Jackson Laboratory (Bar Harbor, ME) and maintained under protocol AABI3617, established by the Institute of Comparative Medicine at Columbia University, in accordance with the Guide for the Care and Use of Laboratory Animals^27^ (institutional animal welfare assurance no. A4711-01). All mice were maintained under SPF conditions on a 12-h light–dark cycle at an ambient temperature of 21.5 ± 1 °C with relative humidity between 30% and 70% and were provided food and water *ad libitum*. Transgenic mice were monitored for onset of disease symptoms, which included hindlimb weakness and weight loss. Lumbar spinal cords were collected from three mice per genotype at P30, P70 and P90 time points. Additional spinal cords were collected from a P30 hSOD1-G93A female mouse and a P64 hSOD1-G93A female mouse. These samples were used for validating antibodies and testing ImmunoSABER repeatability (see **Supplementary Table 3**). Euthanasia was performed using carbon dioxide. Within 5 min of death, mice were transcardially perfused with ice-cold 1× PBS (Gibco, Thermo Fisher Scientific, USA). The spinal cords were then dissected, snap-frozen in ethanol pre-chilled on dry ice, embedded in Optimal Cutting Temperature compound (OCT, Fisher Healthcare, USA) within large molds (VWR, USA), and finally stored at −80 °C until sectioned. Four 10-µm-thick cryosections per animal were collected from nearby levels of the lumbar spinal cord, mounted onto partially assembled PySeq2500 flow cells, and stored at −80 °C for no more than 2 days before processing.

### Flow cell preparation

Untreated glass slides (25 × 75 mm) were drilled with four 0.5-mm-diameter holes (Potomac Photonics, USA) positioned to align with the inlet and outlet ports of the HiSeq 2500 flow-cell stage, and then cleaned in a bath sonicator. The slides were then coated with 0.01% poly-L-lysine (Sigma-Aldrich, P8920) for 5 min followed by overnight drying at room temperature. To form the flow-cell chamber, the flow cell outline was cut from 4 mil (101.6 µm) double-sided polyimide tape (Bertech PPTDE-2, USA) using a Cricut Explore Air 2 (Cricut, 2004419) and attached to the glass slide, defining a 7.75 cm² imaging area that accommodates up to 12 spinal cord sections. Tissue sections were placed into the open flow cell and stored at −80 °C until use. Before starting a PySeq2500 run, the protective film was removed from the double-sided tape and a 25 × 75 mm #1.5 cover glass (Ibidi 10812, Germany) was adhered to the slide to seal the chamber.

### PySeq2500 instrumentation and protocol automation

PySeq2500^18^ provided integrated control of the HiSeq 2500’s three-axis stage, fluidics, temperature control, optics, 532- and 660-nm lasers, and four Hamamatsu TDI line-scan cameras, which capture four emission channels simultaneously (558/32, 610/60, 687/20 and 740/60 nm) through a Nikon 20×/0.75 NA objective. To automate the protocols, recipes were written to pump reagents, incubate flow cells at specified temperatures for specified times, and image premeasured regions of interest (ROIs)^28^. Additionally, configuration files were written to set laser power used for focusing/imaging, flow rates for pumping and other experiment details. Laser power was set to 250 mW, full autofocus routines were used, and reagents were pumped at 100 µL/min. For each experiment run, the configuration files were updated with the bounding boxes specifying the ROIs^28^. An automated experiment was started using the PySeq2500 Python package (v0.64) which confirms experiment settings, primes reagent lines, and then runs the specified protocol without human intervention. The software saves a log of each run and the raw images as 16-bit TIFF files. Because the cameras are line-scan devices, each image is 2,048 pixels wide (the sensor width) and its length is set by the scanned ROI.

### Antibody oligo conjugation

Antibody–oligonucleotide conjugation builds on the CITE-seq antibody-barcoding approach^29^. Here, trans-cyclooctene–tetrazine (TCO–Tz) ligation was performed as described in the CITE-seq Hyper Antibody-Oligo Conjugation protocol^30^. Briefly, 35 nmol of 5′-amine-modified oligomers (IDT, USA) was reacted with 700 nmol TCO-PEG4-NHS (Click Chemistry Tools, CCT-A137-2, USA) in pH 7.6 Borate Buffered Saline (BBS) for 30 min at room temperature and then purified with Micro Bio-Spin P-6 columns (Bio-Rad, 7326221, USA). Then, 50 µg of each antibody was reacted with 10 nmol mTz-PEG4-NHS (Click Chemistry Tools, CCT-1069) in BBS for 30 min at room temperature, washed, and concentrated on 30 kDa cutoff centrifugal filters (EMD Millipore, UFC5003, USA). For each conjugation reaction, 750 pmol of TCO-modified oligonucleotide was incubated with 50 µg of mTz-modified antibody overnight at 4 °C. The reaction was quenched with 7.5 µL TCO-PEG4-GLY for 5 min at room temperature. Conjugation was confirmed by running 1 µL of antibody on a 2% agarose E-Gel (Invitrogen, G401002). Conjugates were stored with 5 µL of 20 mg/mL BSA (NEB, B9200S, USA) and 10 µL of 0.1 M sodium azide (Sigma-Aldrich, 08591-1ML-F, USA) at 4 °C. Conjugates, 10 µL each, were either precipitated all together or individually with 1 mg/mL BSA just prior to staining tissue with 50% saturated ammonium sulfate (∼2.15 M) by incubating on ice for 15 min and then centrifuging at 18,000 g for 15 min at 4 °C in low bind protein tubes (Eppendorf, 05-414-205). Unreacted oligomers were removed by washing 7 times with 1× PBS (Corning, 21-040-CV) up to 500 µL in 50 kDa cutoff centrifugal filters (EMD Millipore, UFC5050, USA), resulting in 40 µL total of mixed antibodies. The following antibodies were conjugated with this method: TOMM40, A17123A; MBP, SMI 2094; ELAVL2, HL1978; MAP2, Poly28225; ALDH1L1, N103/39; AQP4, E5206; SOD1, O98B10; LAMP2, MAC-3; ApoE, 3B3C32; TREM2, 6E9. The rest of the antibodies were received from BioLegend. All antibody information is present in **Supplementary Table 1**.

### Antibody validation

Commercially conjugated antibodies were validated with manual single-plex ImmunoSABER (see *Manual ImmunoSABER*) at 1:100 and 1:200 dilutions. The tissue was counterstained with 50 µL DAPI diluted 1:1000 in 1× PBS for 10 min and then washed with 100 µL PBS before imaging. Nuclei were additionally imaged using a 405 nm laser (LM-HLE-405-1000) with a 445/46 nm bandpass filter (TR-DFLY-F445-046). Antibodies conjugated in house were validated with conventional immunofluorescence before conjugation, after conjugation, and again after conjugation by single-plex ImmunoSABER. For conventional immunofluorescence, spinal cords were sectioned, fixed and stained as described for manual single-plex ImmunoSABER. After primary staining, the sections were washed 3× with 50 µL 1× Intracellular Staining Permeabilization Wash Buffer (BioLegend 421002, USA; hereafter wash buffer) for 10 min at room temperature. Sections were then stained with secondary antibodies diluted 1:200 in the wash buffer for 2 h at room temperature (see **Supplementary Table 1** for secondary antibodies). Sections were washed 3× with 100 µL 1× PBS, and then imaged as in manual single-plex ImmunoSABER. Tiles were stitched using the Andor Fusion integrated image stitcher. Post-conjugation conventional and ImmunoSABER single-plex immunofluorescence images were visually inspected to confirm the staining pattern matched conventional immunofluorescence in expected regions and signal intensity was at least 3× tissue background. Staining patterns were confirmed again with small panels of TotalSeq B, TotalSeq A, TotalSeq D, and our conjugated antibodies by using multiplexed ImmunoSABER (see *Automated ImmunoSABER*). Multiplexed images were processed through the spectral unmixing step using the Hudson image analysis workflow and then visually inspected to confirm the staining pattern matched conventional immunofluorescence as above.

### Concatemer synthesis by primer exchange reaction

Concatemers for ImmunoSABER were made using a primer exchange reaction in 100-µL volumes^15^. Reactions containing 0.3 mM each of dATP, dTTP and dCTP (NEB, N0446S, USA), 6 mM MgSO₄, 2 µL Bst 3.0 DNA polymerase (New England Biolabs, M0374, USA), and 0.2 µM hairpin (5′-CCCCGAAAGTGGCCTCGGGCCTTTTGGCCCGAGGCCACTTTCG-3′) in 1× Isothermal Amplification Buffer (NEB, USA) were mixed on ice and the hairpin oligomer was added to the reaction at the final concentration of 0.5–1 µM. The reaction was heated at 37 °C for 5 min and then hairpin primer was added to a final concentration of 1 µM. Concatemers were generated by extending hairpin primer at 37 °C for 4 h, heat inactivating at 80 °C for 5 min and cooling to 12°C. The concatemers were stored at 4 °C until use. The sequences for the hairpins and primers are listed in **Supplementary Table 2**.

### Tissue preparation

Sections on partially assembled flow cells were thawed at 37 °C for 30 s and then post-fixed with 3.6% formaldehyde (Sigma-Aldrich, F8775) in 1× PBS for 10 min at room temperature. Sections were washed 3 times with 2 mL of 1× PBS for 2 min at room temperature, then permeabilized and blocked simultaneously for 30 min at RT in wash buffer containing 20 nM blocking concatemer (see Supplementary Table 2) and 22 µg/mL mouse IgG Fc fragment (Jackson ImmunoResearch, 015-000-008). Antibodies were diluted in 4 mM EDTA (Invitrogen, AM9849) + 0.2% Dextran Sulfate (bioWORLD, 9011-18-1) with 20 nM blocking concatemer in 1× wash buffer and incubated on the tissue sections at 4 °C overnight (**Supplementary Table 1** for dilutions). The next day, sections were washed 3 times with 1× wash buffer for 10 min at RT. Sections were then post-fixed for 10 min with 3.6% formaldehyde in 1× PBS at room temperature and washed 3 times in 1× PBS for 2 min at RT. Finally, the flow cells were sealed with full-length cover glass (see *Flow cell preparation*) and filled with 0.1% Tween-20 (Sigma-Aldrich, P1379, USA) in 1× Saline-Sodium Phosphate-EDTA (SSPE, Invitrogen, AM9767).

### Manual ImmunoSABER

Manual ImmunoSABER assays were performed with spinal cord sections on poly-L-lysine-coated slides. The tissue was fixed, blocked, permeabilized, stained with antibodies, and post-fixed as described in the previous section. The slide was enclosed in a hybridization cassette (Arrayit, AHC1X24, USA) instead of with a coverslip. A 100 µL volume of concatemer solution containing 25 nM concatemers in 10% dextran sulfate + 30% deionized formamide (ThermoFisher, AM9342, USA) + 0.1% Tween-20 + 2× SSPE was hybridized overnight at 37 °C. Sections were then washed 3× with 50% formamide (EMD Millipore, 4610) in 1× PBS for 10 min at room temperature, followed by 2× washes with 1× PBS at 37 °C for 10 min. Next, repeated cycles of probe hybridization, imaging and stripping were performed, 3 cycles in total. Imagers were hybridized by incubating the sections with 50 µL of a fluorescent DNA imager (250 nM probe, 10 nM blocking concatemer in 1× PBS, **Supplementary Table 2**) at 37 °C for 1 h. Sections were then washed for 5 min with 150 µL of 1× PBS at 37 °C, followed by 2 washes with 150 µL of 1× PBS at 20 °C. The slide was removed from the cassette and tissue was spotted with a few droplets of pH 7.6 700 mM N-acetylcysteine (imaging buffer) before a #1.5 cover glass was applied. Fluorescent images of the sections were acquired with a widefield DragonFly 200 microscope using a Zyla 4.2 MP sCMOS camera with either 0.45 NA 10X (NIKON-MRD00105) or 0.75 NA 20X objectives, the 560 nm laser (LM-HLE-560-1000) with a 594/43 nm bandpass emission filter (TR-DFLY-F594-043) and the 660 nm laser (LM-640-700-DUAL-MM) with a 685/47 (TR-DFLY-F685-047) nm bandpass emission filter to image the Cy3 and Cy5 imagers, respectively. A custom protocol was set up to image a 3 × 3 tile volumetric montage using a 20× 0.75 NA Nikon air objective centered on the central canal of the spinal cords. After imaging, the slide was immersed in 1× PBS and the coverslip was removed. Then the slide was re-enclosed in the hybridization cassette and the fluorescent imagers were stripped from the sections with 3× 150 µL washes in 50% formamide in 1× PBS at 20 °C for 5 min, and 1× 150 µL PBS wash at 37 °C for 10 min.

### Automated ImmunoSABER

Initially, 0.5 mL of concatemers diluted to 25 nM in 10% dextran sulfate + 30% deionized formamide (ThermoFisher, AM9342, USA) + 0.1% Tween-20 + 2× SSPE was pumped through the flow cell and then incubated at 37 °C for 4 h. After the concatemer incubation, the viscous concatemer solution was flushed from the system by pumping 0.5 mL of 1× PBS twice, and then pumping a total of 6 mL of 1× PBS at 20 °C. Next, unbound concatemers were washed 3× with 0.5 mL of 50% formamide in 1× PBS at RT for 10 min, followed by 2 mL of 1× PBS at 37 °C and 2 mL of 1× PBS at 20 °C. The tissue sections were stained with 250 nM of a Cy5-labeled imager complementary to the blocking concatemer, for 1 h at room temperature. After images were acquired (see *Automated image acquisition*), the sections were washed 3× with 0.5 mL of 50% formamide in 1× PBS at RT for 10 min, followed by 2 mL of 1× PBS at 37 °C and 2 mL of 1× PBS at 20 °C. Next, repeated cycles of probe hybridization, imaging and stripping were performed. Briefly, each cycle consisted of incubating 0.5 mL of the fluorescent imager probe set (250 nM each imager probe + 10 nM blocking concatemer in 1× PBS) at 37 °C for 1 h, a 5 min 1× PBS wash at 37 °C, washing with 0.5 mL 1× PBS wash at 37 °C, washing with 2 mL 1× PBS wash at 20 °C, imaging 10 z planes (see *Automated image acquisition*), 3 × 10 min washes in 50% formamide in 1× PBS at 20 °C, and 1× PBS wash at 20 °C. A total of 10 cycles were used for the full 28-antibody panel and 3–5 cycles were used for subpanel validations. An additional final cycle was used to stain and image cellular morphology. This cycle consisted of incubating 0.5 mL of 125 nM Cy3-labeled poly(dT) (IDT) and 8.3 µM DRAQ5 (ThermoFisher, 62251) at 37 °C for 1 h, followed by washing and imaging as above (no stripping).

### Automated image acquisition

Sections were imaged by first filling the flow cells with 0.75 mL imaging buffer (IB, pH 8.0 700 mM N-acetylcysteine, Sigma Aldrich, A7250, USA). If an optimal objective position had not been previously found for the ROI, the “full once” autofocus routine was used to determine the optimal focus position. Briefly, an out-of-focus image of the entire ROI at a fixed objective position was acquired with 1.0 ND filters attenuating the green and red laser power to 25 mW. ROIs within the out-of-focus image were ranked based on contrast, JPEG file size, and relative position. A stack of images at multiple objective positions over iterative ROIs was acquired until a consensus optimal objective focus position was found. The focus position for the ROI was saved and used for subsequent imaging cycles. Stacks of in-focus images at 4.1 µm objective spacing were acquired with 250 mW green and red lasers.

### Manual and automated ImmunoSABER processing repeatability

Mouse spinal cord sections stained with DNA-conjugated anti-GFAP antibody were subjected to repeated hybridization, imaging and stripping of the same fluorescent imager using manual or automated ImmunoSABER protocols. Three manual imaging cycles and five automated imaging cycles were performed. Fluorescence-intensity distributions were assessed in the white matter, ventral horn, central canal and dorsal horn (**Supplementary Fig. 1e**). Signal intensities were arcsinh-normalized on a per-section basis using the tissue median as the background. Differences between sections within an imaging cycle and differences within the same section across imaging cycles were assessed using the two-sample Kolmogorov–Smirnov (KS) statistic (SciPy v1.17.0, `scipy.stats.ks_2samp`). Differences were flagged when the absolute KS statistic was ≥ 0.2 and the 95% bootstrap interval of the signed statistic excluded zero. Bootstrap intervals were estimated from 1,000 resamples, each drawing pixels with replacement from the tested group and its pooled comparison, with each sample containing 10% of the tested group’s pixel count.

### Image analysis workflow

Multicycle fluorescence images were processed using Hudson, an image analysis pipeline implemented in Snakemake^31^. A YAML configuration file specified the experiment directory, imaging-cycle and channel annotations, and processing parameters. The workflow generated processed and spectrally unmixed images, cellular segmentation labels and segment-by-feature matrices, together with a summary report compiled from processing logs. The individual Hudson processing steps are described below.

### Raw image preprocessing

Raw TIFF images were converted into five-dimensional arrays with TCZYX axis ordering and stored in Zarr format. Experiment logs and configuration files were copied to the working directory. Images subsequently underwent sensor-background correction, optional focus projection, channel registration and tile stitching. Sensor-background correction used dark-pixel calibration values obtained during instrument commissioning. For focus projection, JPEG-compressed image sizes were compared across z planes within sliding XY windows. Sharper image patches contain more high-spatial-frequency detail and generally produce larger JPEG files under fixed encoding settings. The z plane yielding the largest compressed file size was selected for each window. The four emission channels were aligned using subpixel translations in X and Y, estimated by phase cross-correlation of images of diffraction-limited TetraSpeck beads (Invitrogen, T7279, USA) acquired during instrument commissioning. Each image strip spanned the full Y extent of the ROI. Strips were stitched along X by removing overlapping pixels according to the experiment configuration file.

### Spectral unmixing

The next step in the pipeline unmixes spectrally overlapping images. Spectral overlap between fluorescence channels was corrected using PICASSOnn, a modified implementation of PICASSO^32^, which estimates spectral spillover by minimizing mutual information between source images and corrected sink images. PICASSOnn introduces two modifications to the original approach. First, we explicitly account for background fluorescence when estimating spectral contamination. For each sink channel, the unmixed image *U* is obtained by subtracting the estimated contributions from spectrally overlapping source channels from the measured image *M*:

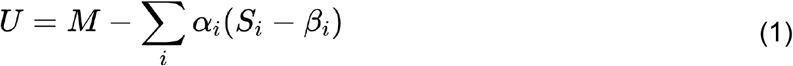

where *S_i_* denotes source image *i*, *α_i_* is its spectral spillover coefficient, and *β_i_* is its estimated background intensity. The subtraction is performed pixel-wise, with each source channel contributing in proportion to its background-subtracted intensity. Second, rather than computing mutual information directly, PICASSOnn uses a neural network-based estimator to enable gradient-based optimization of the spillover coefficients and background parameters (*α_i_*, *β_i_*). These parameters are jointly optimized by stochastic gradient descent using estimated mutual information as the objective. An additional contrast-loss regularization term prevents excessive background subtraction and degenerate solutions in which source signals are suppressed to minimize mutual information. The optimized parameters are subsequently applied to generate spectrally unmixed images, which are stored in CXY OME-NGFF Zarr format, with channel metadata identifying the corresponding protein and fluorophore.

### Cellular segmentation

Cells and cellular segments were identified using Cellpose^33^ based on poly(dT) cytoplasmic and DRAQ5 nuclear channels. Cellular segmentation masks were saved as a TIFF file. Images were segmented using the default tissuenet TN3 model. Using images of mouse spinal cord sections stained with Cy3-labeled poly(dT) and DRAQ5, we fine-tuned a spinal-cord-specific model with the Cellpose Human-in-the-Loop (HITL) training tool. This model was used for all experiments in this study. The custom model was iteratively improved by segmenting 100 training images (512 × 512 px) with the model, manually correcting the segments, and then updating the model. All sections were segmented using summed images of poly(dT) for the cytoplasmic channel and DRAQ5 for the nuclear channel, which resulted in segments of cells and fragmented processes.

### Feature extraction

Morphological features, fluorescence intensities and ViT image embeddings were extracted for each segment and stored in an HDF5-backed MuData object. Segment size, shape and centroid coordinates were measured from the cellular segmentation masks using the `regionprops_table` function in scikit-image v0.22.0^34^. Mean fluorescence intensity was calculated for each protein and dye channel using the same function. A spatial graph was constructed by Delaunay triangulation of segment centroids, with edges filtered using a Euclidean-distance cutoff based on the mean segment diameter. ViT image embeddings were extracted using a pretrained vision transformer (ViT)^35^, specifically `vit_base_patch16_224_miil.in21k`, pretrained on ImageNet-21K-P^36^. Three-channel images were assembled from ELAVL2, GFAP and DRAQ5 signals for crops corresponding to individual segments. Crops were resized, center-cropped and normalized according to the model specifications, then passed through the model to obtain a 768-dimensional ViT image embedding for each segment.

### Image quality control

Each section was manually inspected from overlaid poly(dT) and DRAQ5 images. Folds, tears, and out-of-focus portions of the section were masked off from further analysis.

### Motor neuron identification

Motor neurons were visually identified from overlaid NeuN and poly(dT) images. Large multipolar cells with bright NeuN nuclei located within the ventral horns were marked for further analysis.

### Single-cell embedded latent Dirichlet allocation (sceLDA)

sceLDA combines a conventional variational autoencoder (VAE^37^) with spatial latent Dirichlet allocation (SLDA^21^) in a sequential, two-stage workflow (**Supplementary Fig. 3a**). For each cell *i*, the input consists of a *Q*-dimensional feature vector *v_i_*, a two-dimensional spatial coordinate *r_i_* = (*x_i_*, *y_i_*) and a tissue-section identifier *g_i_*. The VAE first learns a 10-dimensional latent embedding *z_i_* from the cell-level feature vectors using the standard reconstruction and Kullback-Leibler regularization objective. The resulting embeddings are then converted into discrete phenotypic words for spatial topic modeling. The VAE and SLDA components are fitted sequentially rather than through a joint optimization objective; the VAE and SLDA inference algorithms otherwise follow their established formulations.

### Spatially informed phenotypic word construction

The first modification to image-based SLDA concerns the representation of visual words. Rather than applying image convolutions to a regular pixel lattice, we construct a local descriptor for each cell by Gaussian-weighted aggregation of VAE embeddings from cells in the same tissue section. Assuming an isotropic two-dimensional Gaussian kernel, this operation is written as

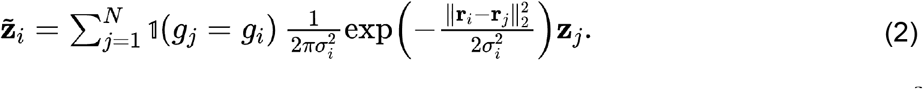

where *N* is the number of cells, 1(*g_j_* = *g_i_*) restricts aggregation to the same section, and 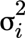 determines the spatial scale of the filter. Thus, all cells within a section may contribute to the descriptor, with contributions decreasing as a function of spatial distance. K-means clustering of the filtered embeddings assigns each cell a phenotypic word label *w_i_* from a codebook of size *V*^38^. Unlike visual words used in image-based SLDA, these labels capture locally aggregated cellular phenotypes rather than image-patch descriptors.

### Spatial document placement and topic inference

The second modification adapts the locations of overlapping spatial documents to the irregular distribution of cells. Whereas image-based SLDA places document centers uniformly across the pixel lattice, sceLDA samples them from observed cell coordinates, making document density reflect the spatial sampling of cells. With phenotypic word assignments fixed, anatomical topic labels and spatial document memberships are inferred using the standard SLDA collapsed Gibbs sampling procedure^21^. Topic updates depend on word-topic and document-topic frequencies, whereas document updates additionally account for topic composition and Gaussian proximity to candidate document centers. Assignments between different tissue sections are excluded. The resulting anatomical topics thus capture spatially recurrent combinations of local phenotypic words rather than whole-section co-occurrence.

### Implementation and hyperparameters

The default VAE encoder contains one hidden layer of 100 units, with batch normalization, ReLU activation, and 20% dropout, followed by a 10-dimensional latent representation obtained through standard VAE reparameterization. The decoder mirrors the encoder and has no final output activation. For spatial topic modeling, we use a phenotypic codebook of *V* = 20 words and set the number of spatial documents to approximately one quarter of the number of cells. The Gaussian variance used for local embedding aggregation is determined from the squared Euclidean distance to the sixth nearest neighboring cell; the analogous spatial scale for each document center is determined from the squared distance to its sixth nearest neighbor. These scales adapt the filtering and document-assignment kernels to the spatial sampling density.

### Synthetic datasets

Synthetic datasets with known spatial topics were generated using three spatial designs: split, checkerboard and polygons (**Supplementary Fig. 3b**). For topic recovery, each method was evaluated on 10 synthetic datasets generated with different random seeds (**Supplementary Fig. 3c–e**). Predicted topic labels were aligned to the true labels by maximum spatial overlap before calculating pixel-wise accuracy and the Jaccard index (see Benchmarking metrics). Runtimes were evaluated separately on synthetic datasets containing 900 to 3,600 cells (**Supplementary Fig. 3f**).

### Related methods

In addition to building the VAE and custom SLDA algorithm for sceLDA, we created wrappers for existing NSF^39^, NSFH^39^, and MEFISTO^40^ (v0.7.2) implementations to facilitate fair testing and comparison^39,40^. To do this, each wrapper converts raw array data with embedded image indices and spatial coordinates into AnnData objects consistent with the format expected by each method. Additionally, we compared against the LDA, NMF^41^, and K-Means implementations provided by the scikit-learn Python library^42^ (v1.5.0 `sklearn.decomposition.LatentDirichletAllocation`, `sklearn.decomposition.NMF`, and `sklearn.cluster.KMeans`, respectively) with default parameters other than random initializations for NMF and K-Means. We configured all workflows with 500 inducing points and used those in all comparisons.

### Benchmarking metrics

We used pixel-wise accuracy, the weighted Jaccard index and the Calinski–Harabasz^43^ index (v1.5.0, `sklearn.metrics.accuracy_score`, `sklearn.metrics.jaccard_score`, `sklearn.metrics.calinski_harabasz_score`) to evaluate the performance of spatial topic modeling methods on spatial datasets in instances where ground truth was available. Pixel-wise accuracy measured the proportion of identical cell or pixel topic labels between two spatial topic modeling results after aligning their labeling schemes through maximum spatial overlap. The weighted Jaccard index summarized overlap between inferred and true topic assignments. The Calinski–Harabasz index was used to quantify cluster distinctness by taking the ratio of between-cluster separation to within-cluster dispersion across predicted topic assignments. Metrics were summarized across the 10 synthetic datasets generated with different random seeds.

### Motor neuron measurements

For each segment and channel, the mean raw intensity was calculated first and then transformed as ln(1 + mean intensity) to obtain the segment signal. For each section and channel, the median signal of motor-neuron segments was normalized by subtracting the median signal of all cell-sized, non-motor-neuron segments inside the cord in the same section (**Fig. 2d–f**). Signals in the 50-µm neighborhoods (**Fig. 2e,f**) were measured as the median signal of cell-sized, non-motor-neuron segments whose centroids lay within 50 µm of a motor-neuron centroid (SciPy v1.18.1, scipy.spatial.cKDTree), normalized to the same section reference. For display in **Fig. 2e,f**, mouse-level normalized signals were exponentiated after subtracting the mean normalized log signal of wild-type mice at the same age. Motor-neuron counts per ventral horn were analyzed on the log scale (**Fig. 2c**). The p values of **Fig. 2d** were adjusted over 90 tests (30 channels × 3 ages) with the Benjamini–Hochberg procedure^44^ (statsmodels v0.15.0, `statsmodels.stats.multitest.multipletests`).

### Region masks

We generated per-section anatomical region masks by warping a single region map based on the 11 anatomical regions defined by Barretto et al.^11^ (dorsal horn, medial gray, central canal, ventral horn, dorsal-medial, medial-lateral, ventral-lateral and ventral-medial white matter, and dorsal, lateral and ventral edges) back into each section’s native image frame (see *Spatial topic probability maps*). To separate gray from white matter, cells were first z-scored per section using ELAVL2 + NeuN − MBP expression + local segment area (the mean area of all segments within 50 µm). The cell scores were pooled and kernel-smoothed on the template grid (Gaussian, σ = 150 template px; 50-px bins). Bins were classified as gray matter when a two-component Gaussian mixture gave a posterior probability of gray matter above 0.5. Gray matter was split by height into dorsal horn (top quarter of cord), ventral horn (below the canal) and medial gray (the remaining gray matter). Each interior white-matter bin was assigned the label of the gray-matter face nearest to it. White matter in the outer 10% of the cord radius was labeled as the edge region next to its white-matter sector. Gray-matter pixels within 75 µm of the labeled central canal were relabeled as central canal. These template masks were then manually corrected for each section.

### Topic annotation and marker profiles

The marker profile of a topic was computed per section as the median signal of the topic segments minus the median of all segments of the section, averaging over the sections of a mouse and then over the 18 mice (**Fig. 3b**). Topics were grouped by the Euclidean distance between the mean profiles of two topics (30 channels, log2; mice in which both topics were present), divided by the root-mean-square between-mouse spread of a topic profile. Topics were clustered by complete linkage on this ratio (`scipy.cluster.hierarchy.linkage`), and the tree was cut at the number of clusters that remained stable over the widest range of cut heights (six; 4–9 considered). The two non-reactive gray-matter topics, T4 and T13, fell into separate clusters but are shown as one family. The P90 run, whose sections contained a higher proportion of small segments (53% below 40 µm^2^, against 13% at P30 and 23% at P70), assigned to T13 most of the non-reactive gray matter that the P30 and P70 runs assigned to T4.

### Spatial topic probability maps

To compare the spatial distributions of topics across sections (**Fig. 3c**), each section was registered to a common template using five hand-placed landmarks: the central canal, the dorsal median septum, the ventral median fissure, and the left and right lateral edges. Sections were rotated dorsal-side up with the origin at the central canal on the midline. Each quadrant was then scaled by the section’s own half-width and dorsal/ventral extent to the means over the 70 sections. For each section, in-cord segments were binned at 30 µm, left and right sides were averaged, and counts were smoothed with a Gaussian kernel (s.d. 45 µm; `scipy.ndimage.gaussian_filter`). The share of each topic among all segments was then averaged over the sections of a mouse and over the mice. The outline encloses positions covered by at least half of the sections.

### Motor neuron neighborhoods and regional topic shares

GFAP-high neighborhoods were defined by the two GFAP-high topics of the ventral horn, T14 and T9. Non-reactive gray matter included T4 and T13. The neighborhood of a motor neuron comprised the cell-sized, non-motor-neuron segments whose centroid lay within 50 µm of its centroid. A motor neuron was counted as embedded in a GFAP-high neighborhood when T14 or T9 was the most frequent topic among them (**Fig. 4a,b**). Zones around motor neurons (**Fig. 4c**) consisted of the cell-sized, non-motor-neuron segments inside the cord within a distance r of any motor-neuron centroid (r = 25, 50, 75, 100, 150 and 200 µm; each zone contains the smaller ones). Regional shares (**Fig. 4d**) were computed among all segments of the ventral horn and among white-matter segments.

### Spatial pooling of GFAP

Cell-sized segments (area ≥ 80 µm²) were pooled into circular spots of 55 or 100 µm diameter on fixed grids with 100 or 200 µm center-to-center spacing, respectively, anchored at the image-coordinate origin. Spot GFAP was the area-weighted mean raw intensity of segments whose centroids lay inside the spot. Spots with fewer than three segments were excluded, and ventral-horn spots required more than half their segments to lie in the ventral horn. At every scale, the section-level value was ln(1 + median ventral-horn intensity) – ln(1 + median whole-section intensity), using only segments or spots inside the cord. Values were analyzed with the slide-balanced mouse-level model (Statistical analysis; Supplementary Fig. 4).

### Comparison with spatial transcriptomics

Module maps of Maniatis et al.^5^ (Figs. S6 and S8, group means of SOD1-G93A and SOD1-WT mice at P30, P70 and P100) were digitized by K.P. They were then placed on a normalized half-cord frame given by the tissue bounding box of each panel, averaged over left and right, and subtracted to obtain SOD1-G93A minus SOD1-WT. Our topic maps were computed as in Spatial topic probability maps, summarized per group as the median of the three mice, and differenced as SOD1-G93A minus wild-type. Both were compared on a shared 10 × 16 half-cord grid, matching their P100 to our P90. Pearson’s r (NumPy v2.5.3, `numpy.corrcoef`) was computed over all shared bins and over bins in which most segments lay in the ventral horn. Neighboring bins are not independent, so r is descriptive. Differences are drawn in full color where all three SOD1-G93A mice lay above or below all three wild-type mice. For the human comparison (**Fig. 4f,g**), we used the log2 fold changes between ALS and control donors reported by Barretto et al.^11^ for the neuronal module M11 and the GFAP-astrocyte module M9 in 7 regions of post-mortem lumbar spinal cord, read from their figure by K.P. Our region masks follow the same layout, so regions were matched one to one. For each region, the share of all segments assigned to T13 or T14 was compared between genotypes as a log2 odds ratio (see Statistical analysis).

### Statistical analysis

Mice were the unit of analysis (n = 3 per genotype and age). Each age was imaged in a separate run on two slides, each slide holding sections of both genotypes. For each measure (for example, the share of ventral-horn segments assigned to T14), we first computed one value per section. We then removed slide differences by subtracting a slide offset from every section on that slide. The offset was the average of the slide’s wild-type mean and its SOD1-G93A mean. This removes differences between slides but leaves the difference between genotypes within each slide unchanged. Values were averaged per mouse and fitted by ordinary least squares, y ∼ age + age:genotype, over the 18 mice (statsmodels v0.15.0, `statsmodels.formula.api.ols`). The t tests used 9 degrees of freedom: 12 residual degrees of freedom minus the 3 slide offsets. Intensities and counts were analyzed on the log scale and are reported as SOD1-G93A/wild-type ratios. Topic shares are reported as differences in percentage points (**Fig. 4b–d**) or as odds ratios (log2 in **Fig. 4f,g**). T13, assigned almost only in the P90 run, was fitted at P90 alone (3 degrees of freedom). We report 95% confidence intervals and two-sided p values, adjusted for multiple testing only where stated (**Fig. 2d; Supplementary Fig. 3c–e**). For the synthetic-data benchmark (**Supplementary Fig. 3c–e**), each dataset was one observation. sceLDA was compared with each other method using two-sided t-tests, and p values were adjusted with the Benjamini–Hochberg procedure^44^ over the seven comparisons within each metric. Analyses were run in Python v3.12–3.13 with NumPy v2.5.3–2.5.4, pandas v3.0.5–3.1.1, SciPy v1.18.1, statsmodels v0.15.0 and scikit-image v0.26.0, and figures were drawn with Matplotlib v3.11.2.

## Code availability

All the code has been made available at: https://github.com/nygctech/PySeqImmunoSABER

## Data availability

All the data has been made available at: https://doi.org/10.5281/zenodo.23959524

## Acknowledgements

This work was supported by the Knut and Alice Wallenberg Foundation, Beijer Laboratory for Gene and Neuro Research, Swedish Society for Medical Research, MacMillan Center for the Non-Coding Genome, Science for Life Laboratory, and National Institutes of Health (NIH) grants 1U54AG076040-01 and 1RM1HG011014-01 (S.V.); Target ALS grant FD-2023-GEN-S1 (S.V. and H.P.); NIH grants R01NS118183, R01NS116350, R01NS127186, and R01NS118570 (H.P.); Chan Zuckerberg Initiative grant DAF2021-240116 (S.M. and K.P.); and NIH grant 1R01CA272891-01 (S.M.). S.V. is additionally supported as a Wallenberg Academy Fellow and SciLifeLab Fellow at Uppsala University.

## Author contributions

K.P., H.P. and S.V. conceptualized the project; H.P. provided animal tissues for this study; K.P., S.R. and J.S. developed the image processing pipeline with supervision from S.V.; C.F. and S.R. developed sceLDA with supervision from B.D. and S.V.; K.P., Si.M. and K.D. conducted the experiments; Se.M. performed HITL annotation and training; K.P. and M.P. analyzed the data and supported data, software, and scientific project management with guidance from S.V. and help from D.D.; K.P., S.R., M.P., and S.V. interpreted the results; K.P., M.P., and S.V. wrote the manuscript and prepared the figures; all authors discussed the results.

## Supplementary Tables

**Supplementary Table 1.** Early mouse ALS antibody panel and conjugated DNA sequences

**Supplementary Table 2.** Primer, hairpin and imager DNA sequences

**Supplementary Table 3.** Animal age, genotype and sex

---

**Supplementary Figure 1:**
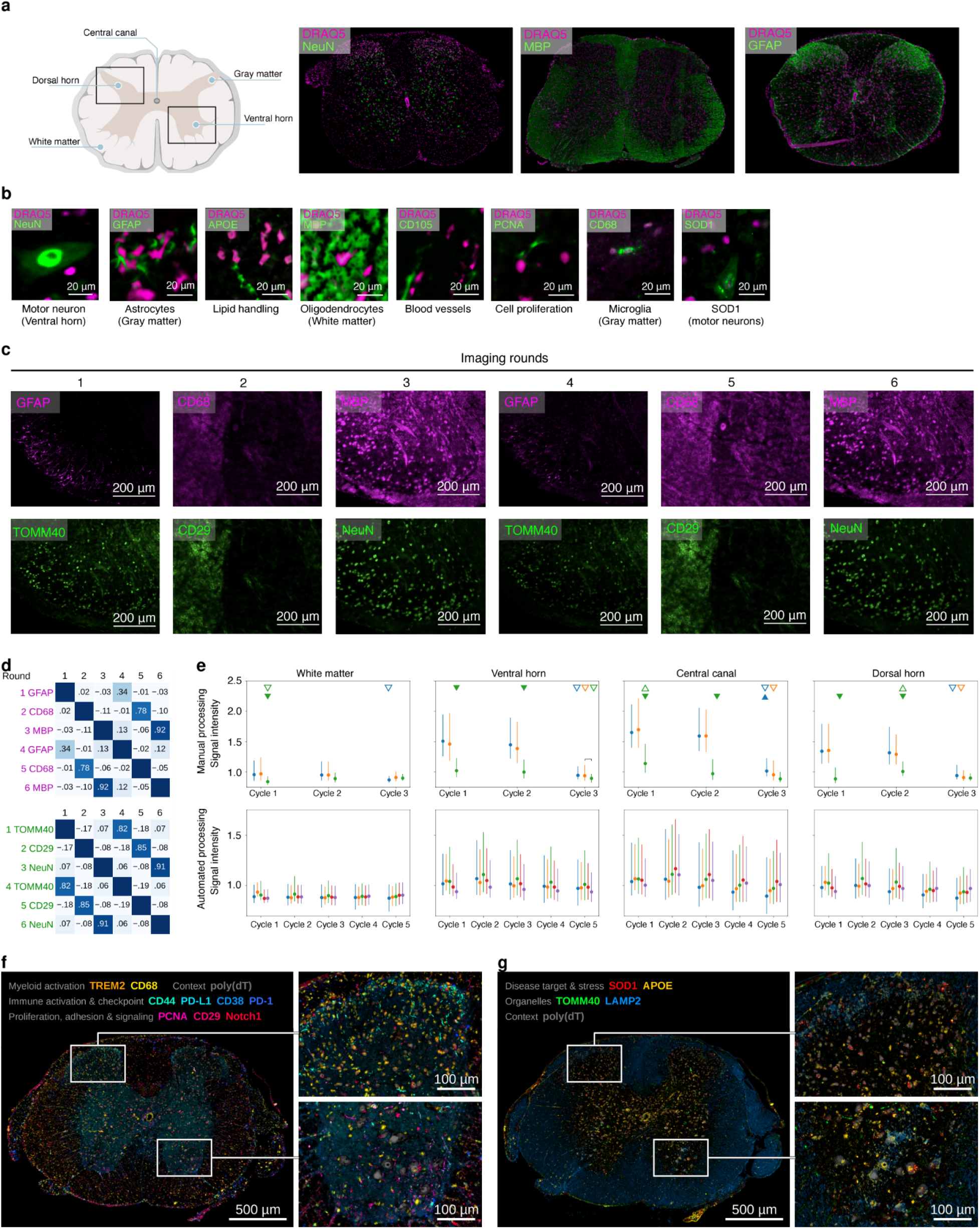
Antibody staining, cyclic imaging repeatability and additional protein-marker images. **(a) Whole-section antibody staining:** Schematic of a transverse spinal cord section (left) and representative fluorescence images of lumbar spinal cord sections stained for NeuN, MBP or GFAP (green), with DRAQ5 nuclear counterstain (magenta). **(b) Antibody staining in selected spinal cord regions:** Cropped fluorescence images showing the indicated markers (green) and DRAQ5 (magenta). Scale bars, 20 µm. Images in a and b were acquired during individual antibody screening, before the sequential multiplexed readout, in spinal cord sections from a P64 female SOD1-G93A mouse. **(c) Repeated detection across imaging cycles:** One field of a mouse spinal cord section imaged over six successive imaging cycles in two channels: GFAP and TOMM40 in imaging cycles 1 and 4, CD68 and CD29 in imaging cycles 2 and 5, and MBP and NeuN in imaging cycles 3 and 6 (magenta and green). Scale bars, 200 µm. **(d) Imaging-cycle repeatability:** Pixel-wise Pearson correlation between every pair of imaging cycles of each channel in c (top, magenta; bottom, green). The same target read three imaging cycles later correlated with its first readout. Different targets in consecutive imaging cycles did not correlate, showing no detectable carry-over from one imaging cycle to the next. **(e) Manual and automated processing:** Signal intensity in the white matter, ventral horn, central canal and dorsal horn over successive imaging cycles of manual processing (top; three imaging cycles) and automated processing on PySeq2500 (bottom; five imaging cycles). Significant shifts between sections within an imaging cycle are denoted by a closed triangle. Significant shifts of the same section between imaging cycles are denoted by an open triangle. Direction of the shift is denoted by the orientation of the triangle. Shifts were tested with a bootstrapped two-sided two-sample Kolmogorov–Smirnov test, with shifts called when the 95% interval of the signed statistic excluded zero and its absolute value was ≥ 0.2. **(f) Activation state:** Composite image of nine activation-state markers (TREM2, CD68, CD44, PD-L1, CD38, PD-1, PCNA, CD29 and Notch1) in the lumbar spinal cord section shown in Fig. 1h (P90 SOD1-G93A mouse), with the dorsal horn (top) and ventral horn (bottom) regions enlarged on the right. Poly(dT) is shown in gray. **(g) Pathology:** Composite image of four pathology markers (SOD1, APOE, TOMM40 and LAMP2) in the same section and regions. Poly(dT) is shown in gray. Marker colors in f and g are indicated at the top of each panel. Scale bars in f and g, 500 µm (whole section) and 100 µm (enlarged regions).

**Supplementary Figure 2:**
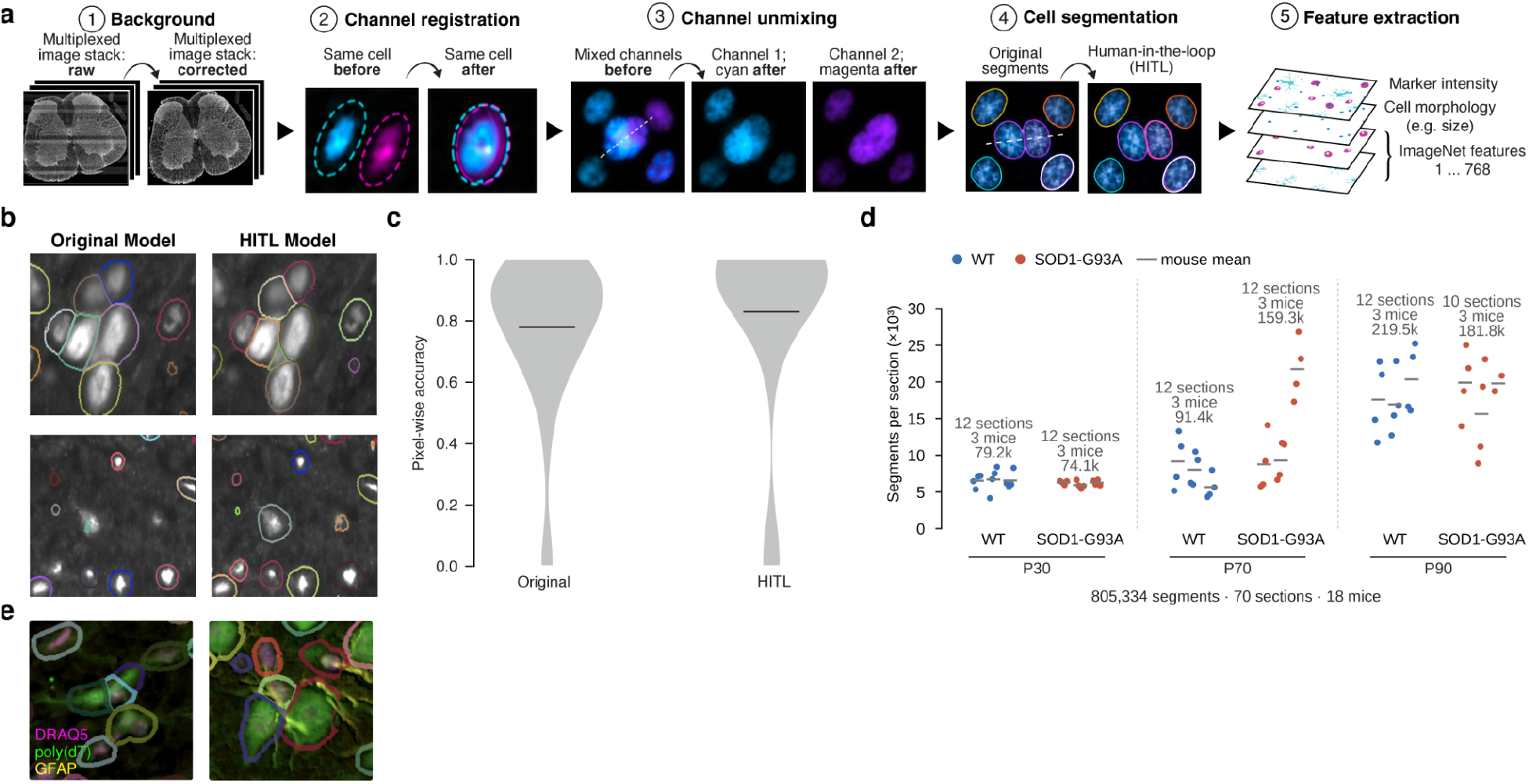
Hudson image processing and cellular segmentation. **(a) Hudson image-analysis pipeline:** Schematic of the five processing steps that convert multicycle fluorescence images into spatially resolved cellular profiles: background correction, channel registration, channel unmixing, cellular segmentation and feature extraction. Extracted features comprise protein-marker intensities, 15 morphological measurements and 768-dimensional ViT image embeddings from a network pretrained on ImageNet. **(b) Cellular segmentation model refinement:** Representative images with segmentation boundaries from the original Cellpose model (left) and the model refined by human-in-the-loop (HITL) training (right). Colored outlines indicate individual segments. Cellular segmentation used poly(dT) as the cytoplasmic channel and DRAQ5 as the nuclear channel. **(c) Cellular segmentation accuracy:** Distributions of pixel-wise accuracy of the original and HITL-refined models, evaluated against reference segmentations. Horizontal black lines indicate median accuracy values. **(d) Cellular segmentation across the tissue cohort:** Number of segments per section in wild-type (blue) and SOD1-G93A (red) mice at P30, P70 and P90. Dots, sections, arranged by mouse; gray lines, mouse means. Numbers above each group give the sections, mice and total segments. In total, Hudson produced 805,334 segments in 70 spinal cord sections from 18 mice. **(e) Examples of cellular segmentation:** Segmentation boundaries overlaid on DRAQ5 (magenta), poly(dT) (green) and GFAP (yellow). Segments include cell bodies and cellular processes, which may be recovered as separate segments.

**Supplementary Figure 3:**
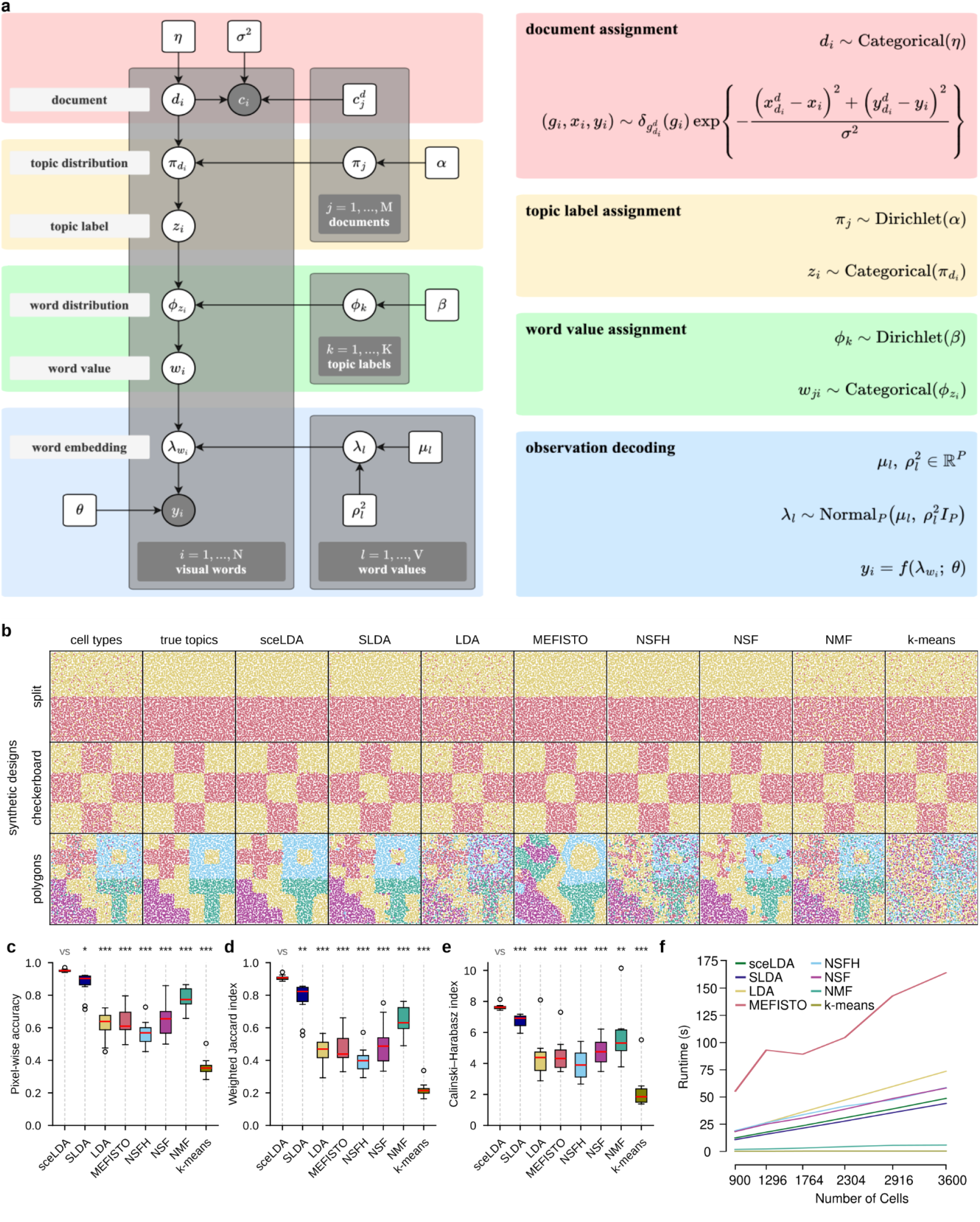
The sceLDA model and its benchmark on synthetic data. **(a) Generative model of sceLDA:** Plate diagram (left) and the corresponding probability distributions (right). Each segment (visual word) is assigned to a spatial document according to its location, to a topic according to the topic distribution of that document, and to a phenotypic word value according to the word distribution of that topic. Word values carry latent embeddings that are decoded into the observed features. Shaded nodes, observed variables; open nodes, latent variables and parameters. Plates denote repetition over segments (N), spatial documents (M), topics (K) and word values (V). **(b) Synthetic datasets:** Three spatial designs (split, checkerboard and polygons). The first two columns show the simulated cell types and the true topics; the other columns show the topics recovered by sceLDA, spatial LDA (SLDA), latent Dirichlet allocation (LDA), MEFISTO, hybrid nonnegative spatial factorization (NSFH), nonnegative spatial factorization (NSF), nonnegative matrix factorization (NMF) and k-means. **(c–e) Topic recovery:** Agreement with the true topics, as pixel-wise accuracy (c) and weighted Jaccard index (d), and separation of the inferred topics, as Calinski–Harabasz index (e), for each method over 10 synthetic datasets generated with different random seeds; higher values indicate better performance. Boxes, interquartile range; red lines, medians; whiskers, the most extreme values within 1.5× the interquartile range; circles, values beyond. Stars, sceLDA vs each method, two-sided t-test with Benjamini–Hochberg correction over the seven comparisons of each metric: *q < 0.05, **q < 0.01, ***q < 0.001. **(f) Runtime:** Execution time of each method against the number of cells in the synthetic dataset.

**Supplementary Figure 4:**
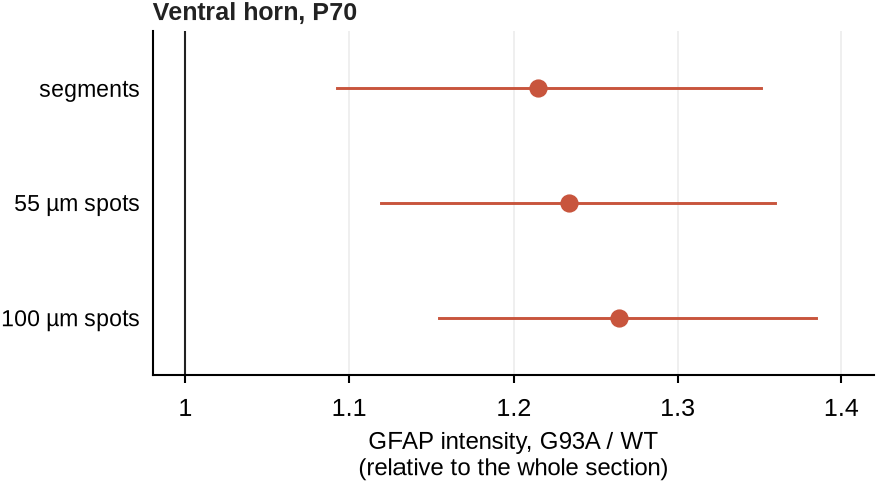
Ventral-horn GFAP measurements at cellular and pooled spatial scales. GFAP intensity in the ventral horn at P70 in SOD1-G93A mice relative to wild type, after normalization to the whole section. Values are shown for cell-sized segments and after pooling into 55-µm spots on a 100-µm grid or 100-µm spots on a 200-µm grid. Dots, estimates; lines, 95% confidence intervals. The same section-level summary and normalization were used at all scales (Methods). n = 3 mice per genotype and age, with mice as the unit of analysis.

## REFERENCES

1. Baccin, C. et al. Combined single-cell and spatial transcriptomics reveal the molecular, cellular and spatial bone marrow niche organization. Nat. Cell Biol. 22, 38–48 (2020).

2. Schürch, C. M. et al. Coordinated cellular neighborhoods orchestrate antitumoral immunity at the colorectal cancer invasive front. Cell 182, 1341–1359.e19 (2020).

3. Keren, L. et al. A structured tumor-immune microenvironment in triple negative breast cancer revealed by multiplexed ion beam imaging. Cell 174, 1373–1387.e19 (2018).

4. Jackson, H. W. et al. The single-cell pathology landscape of breast cancer. Nature 578, 615–620 (2020).

5. Maniatis, S. et al. Spatiotemporal dynamics of molecular pathology in amyotrophic lateral sclerosis. Science 364, 89–93 (2019).

6. Boillée, S. et al. Onset and progression in inherited ALS determined by motor neurons and microglia. Science 312, 1389–1392 (2006).

7. Yamanaka, K. et al. Astrocytes as determinants of disease progression in inherited amyotrophic lateral sclerosis. Nat. Neurosci. 11, 251–253 (2008).

8. Beers, D. R., Henkel, J. S., Zhao, W., Wang, J. & Appel, S. H. CD4+ T cells support glial neuroprotection, slow disease progression, and modify glial morphology in an animal model of inherited ALS. Proc. Natl Acad. Sci. USA 105, 15558–15563 (2008).

9. Kang, S. H. et al. Degeneration and impaired regeneration of gray matter oligodendrocytes in amyotrophic lateral sclerosis. Nat. Neurosci. 16, 571–579 (2013).

10. Zhang, Z. et al. Integrated single-cell and spatial transcriptomic profiling in ALS uncovers peripheral-to-central immune infiltration and reprogramming. Nat. Neurosci. 29, 1735–1748 (2026).

11. Barretto, N., et al. A multi-omics characterization reveals distinct molecular signatures in the human motor cortex and lumbar spinal cord in ALS. bioRxiv (2026) doi:10.64898/2026.07.24.740361.

12. Giesen, C. et al. Highly multiplexed imaging of tumor tissues with subcellular resolution by mass cytometry. Nat. Methods 11, 417–422 (2014).

13. Lin, J.-R. et al. Highly multiplexed immunofluorescence imaging of human tissues and tumors using t-CyCIF and conventional optical microscopes. eLife 7, e31657 (2018).

14. Goltsev, Y. et al. Deep profiling of mouse splenic architecture with CODEX multiplexed imaging. Cell 174, 968–981.e15 (2018).

15. Saka, S. K. et al. Immuno-SABER enables highly multiplexed and amplified protein imaging in tissues. Nat. Biotechnol. 37, 1080–1090 (2019).

16. Rivest, F. et al. Fully automated sequential immunofluorescence (seqIF) for hyperplex spatial proteomics. Sci. Rep. 13, 16994 (2023).

17. Hickey, J. W. et al. Spatial mapping of protein composition and tissue organization: a primer for multiplexed antibody-based imaging. Nat. Methods 19, 284–295 (2022).

18. Pandit, K. et al. An open source toolkit for repurposing Illumina sequencing systems as versatile fluidics and imaging platforms. Sci. Rep. 12, 5081 (2022).

19. Gurney, M. E. et al. Motor neuron degeneration in mice that express a human Cu,Zn superoxide dismutase mutation. Science 264, 1772–1775 (1994).

20. Blei, D. M., Ng, A. Y. & Jordan, M. I. Latent Dirichlet allocation. J. Mach. Learn. Res. 3, 993–1022 (2003).

21. Wang, X. & Grimson, E. Spatial latent Dirichlet allocation. in Advances in Neural Information Processing Systems vol. 20 1577–1584 (2007).

22. Baker, D. J. et al. Lysosomal and phagocytic activity is increased in astrocytes during disease progression in the SOD1 (G93A) mouse model of amyotrophic lateral sclerosis. Front. Cell. Neurosci. 9, 410 (2015).

23. Hall, E. D., Oostveen, J. A. & Gurney, M. E. Relationship of microglial and astrocytic activation to disease onset and progression in a transgenic model of familial ALS. Glia 23, 249–256 (1998).

24. Fischer, L. R. et al. Amyotrophic lateral sclerosis is a distal axonopathy: evidence in mice and man. Exp. Neurol. 185, 232–240 (2004).

25. Gautier, O. et al. An emergent disease-associated motor neuron state precedes cell death in ALS. Cell 189, 5044–5064.e12 (2026).

26. Kelley, K. W. et al. Kir4.1-dependent astrocyte-fast motor neuron interactions are required for peak strength. Neuron 98, 306–319.e7 (2018).

27. National Research Council. Guide for the Care and Use of Laboratory Animals. (National Academies Press, Washington, DC, 1996).

28. Pandit, K., et al. Spatial proteomics on repurposed sequencers reveals early astrocytic remodeling in an ALS mouse model. Zenodo 10.5281/ZENODO.23959524 (2026).

29. Stoeckius, M. et al. Simultaneous epitope and transcriptome measurement in single cells. Nat. Methods 14, 865–868 (2017).

30. Houck-Loomis, B., Stoeckius, M. & Hao, S. CITE-seq Hyper Antibody-Oligo Conjugation. https://citeseq.wordpress.com/wp-content/uploads/2019/03/cite-seq_hyper_conjugation_190321.pdf (2019).

31. Mölder, F. et al. Sustainable data analysis with Snakemake. F1000Res. 10, 33 (2021).

32. Seo, J. et al. PICASSO allows ultra-multiplexed fluorescence imaging of spatially overlapping proteins without reference spectra measurements. Nat. Commun. 13, 2475 (2022).

33. Pachitariu, M. & Stringer, C. Cellpose 2.0: how to train your own model. Nat. Methods 19, 1634–1641 (2022).

34. van der Walt, S. et al. scikit-image: image processing in Python. PeerJ 2, e453 (2014).

35. Dosovitskiy, A., et al. An image is worth 16×16 words: transformers for image recognition at scale. in International Conference on Learning Representations (2021).

36. Ridnik, T., Ben-Baruch, E., Noy, A. & Zelnik-Manor, L. ImageNet-21K pretraining for the masses. in Proceedings of the Neural Information Processing Systems Track on Datasets and Benchmarks vol. 1 (2021).

37. Kingma, D. P. & Welling, M. Auto-encoding variational Bayes. in International Conference on Learning Representations (2014).

38. Winn, J., Criminisi, A. & Minka, T. Object categorization by learned universal visual dictionary. in Tenth IEEE International Conference on Computer Vision vol. 2 1800–1807 (2005).

39. Townes, F. W. & Engelhardt, B. E. Nonnegative spatial factorization applied to spatial genomics. Nat. Methods 20, 229–238 (2023).

40. Velten, B. et al. Identifying temporal and spatial patterns of variation from multimodal data using MEFISTO. Nat. Methods 19, 179–186 (2022).

41. Lee, D. D. & Seung, H. S. Learning the parts of objects by non-negative matrix factorization. Nature 401, 788–791 (1999).

42. Pedregosa, F. et al. Scikit-learn: machine learning in Python. J. Mach. Learn. Res. 12, 2825–2830 (2011).

43. Caliński, T. & Harabasz, J. A dendrite method for cluster analysis. Commun. Stat. 3, 1–27 (1974).

44. Benjamini, Y. & Hochberg, Y. Controlling the false discovery rate: a practical and powerful approach to multiple testing. J. R. Stat. Soc. B 57, 289–300 (1995).

